# Persister Cells Resuscitate Using Membrane Sensors that Activate Chemotaxis, Lower cAMP Levels, and Revive Ribosomes

**DOI:** 10.1101/486985

**Authors:** Ryota Yamasaki, Sooyeon Song, Michael J. Benedik, Thomas K. Wood

**Affiliations:** Department of Chemical Engineering, Pennsylvania State University, University Park, Pennsylvania, 16802-4400, USA; Department of Biochemistry and Molecular Biology, Pennsylvania State University, University Park, Pennsylvania, 16802-4400, USA; Department of Biology, Texas A & M University, College Station, Texas, 77843-3122

**Keywords:** persisters, alanine, glucose, resuscitation, ribosomes

## Abstract

Persistence, the stress-tolerant state, is arguably the most vital phenotype since nearly all cells experience nutrient stress, which causes a sub-population to become dormant. However, how persister cells wake to reconstitute infections is not understood well. Here, using single-cell observations, we determined that *Escherichia coli* persister cells resuscitate primarily when presented with specific carbon sources, rather than spontaneously. In addition, we found that the mechanism of persister cell waking is through sensing nutrients by chemotaxis and phosphotransferase membrane proteins. Furthermore, nutrient transport reduces the level of secondary messenger cAMP through enzyme IIA; this reduction in cAMP levels leads to ribosome resuscitation and rescue. Resuscitating cells also immediately commence chemotaxis toward nutrients, although flagellar motion is not required for waking. Hence, persister cells wake by perceiving nutrients via membrane receptors which relay the signal to ribosomes via the secondary messenger cAMP, and persisters wake and utilize chemotaxis to acquire nutrients.

## INTRODUCTION

The persister cell phenotype was first noted by Hobby *et. al.* in 1942 (Hobby et al., 1942) when penicillin did not completely sterilize a *Staphylococcus aureus* culture (1% of the population remained intact). The surviving subpopulation was deemed ‘persisters’ by Bigger *et. al* in 1944 (Bigger, 1944). Both groups determined that persisters are dormant (Bigger, 1944; Hobby et al., 1942), which has been corroborated (Kwan et al., 2013; Shah et al., 2006), and further research has demonstrated persister cells are not mutants (Chowdhury et al., 2016b; Kwan et al., 2015a) but instead acquire their antibiotic tolerance through this dormancy.

The persister cell phenotype is ubiquitous and has been well described in many bacteria such as *Escherichia coli* (Fisher et al., 2017), *Pseudomonas aeruginosa* (Fisher et al., 2017), *S. aureus* (Fisher et al., 2017) and in Archaea (Megaw and Gilmore, 2017). Critically, the persister state arises not only after antibiotic stress but nutrient stress also creates persister cells (Bernier et al., 2013; Maisonneuve and Gerdes, 2014; Martins et al., 2018); in fact, the classic viable but not culturable state appears to be the same as the persister state (Kim et al., 2018a) so persisters form everywhere as all bacterial cells eventually face nutrient stress (Song and Wood, 2018). Hence, it may be argued that the persister state is one of the most fundamental bacterial phenotypes.

Persister cells form from a reduction in metabolism usually from activation of a toxin of a toxin/antitoxin system. Evidence of this is that the deletion of several toxins of toxin/antitoxin systems such as MqsR (Kim and Wood, 2010; Luidalepp et al., 2011), TisB (Dörr et al., 2010), and YafQ (Harrison et al., 2009) leads to a reduction in persistence. Similarly, production of toxins unrelated to toxin/antitoxin systems can also increase persistence (Chowdhury et al., 2016a).

We have found persister cells resuscitate as soon as instantaneously in rich medium (Kim et al., 2018b) and wake based on their ribosome content (Kim et al., 2018b). For example, persister cells with four fold fewer ribosomes are delayed by several hours in their resuscitation while ribosome levels increase (Kim et al., 2018b). Others have suggested, but not shown, that cells may be resuscitated by reversing the effects of toxins of toxin/antitoxin systems (Cheverton et al., 2016). Hence, it is still unknown what pathway is involved in persister cell waking in regard to nutrient sensing.

To study persister cells without introducing traits of more prevalent cell phenotypes (e.g., slow-growing, tolerant stationary cells), their concentration needs to be increased so they are the dominant phenotype. Previously, we showed how to create a high percentage of *E. coli* persister cells (up to 70%) via rifampicin-pretreatment to stop transcription, carbonyl cyanide *m*-chlorophenylhydrazone (CCCP) to stop ATP production, or tetracycline to stop translation (Kwan et al., 2013); these cells were shown to be *bona fide* persister cells via eight different assays (multi-drug tolerance, immediate change from persistence to non-persistence in the presence of nutrients, dormancy based on lack of cell division in the absence of nutrients, dormancy via metabolic staining and cell sorting, no change in MIC compared to exponential cells, no resistance phenotype, similar morphology to ampicillin-induced persisters, and similar resuscitation as ampicillin-induced persisters) (Kim et al., 2018b). Several other research groups have used our methods to make persister cells (Cui et al., 2018; Grassi et al., 2017; Narayanaswamy et al., 2018; Sulaiman et al., 2018; Tkhilaishvili et al., 2018); for example, Cui *et. al.* used rifampin and tetracycline to induce persistence for screening the *E. coli* Keio library and found several DNA repair mutants (e.g., *recA*, *recC*, *ruvA*, and *uvrD*) with reduced persistence, Grassi *et. al*.(Grassi et al., 2017) used CCCP to generate *P. aeruginosa* and *S. aureus* persister cells to determine that antibiotic-tolerant phenotypes depend on different classes of antibiotics, and Narayanaswamy *et. al.*(Narayanaswamy et al., 2018) used CCCP to generate *P. aeruginosa* persister cells and found poly (acetyl, arginyl) glucosamine is effective for killing them.

In the present study, we found that rather than waking spontaneously, persister cells wake primarily as a result of sensing nutrients. We then elucidated the waking pathway and found persister cells wake via chemotaxis and transport pathways that serve to reduce cAMP, activate ribosomes, and commence chemotaxis.

## RESULTS

### Overview

To create a large enough population of persister cells so that we could investigate resuscitation at the single-cell level, we pre-treated exponential cells with rifampicin to stop transcription since this method converts the rare persister phenotype into the dominant cell state; i.e., we increased the number of persister cells by 10^5^-fold which enables us to study more readily persister cells at the single-cell level (Kwan et al., 2013). The persister cells induced by rifampicin are isolated from any remaining non-persister cells by ampicillin treatment since the non-persisters cells lyse so that the remaining population consists solely of only persister cells (Kwan et al., 2013).

To study these persister cells, we utilized initially agar pates. We reasoned that if specific persister cells wake faster, larger colonies would form on M9 agar plates since we found previously that once they wake, persister cells grow at the same rate as exponentially-growing cells (Kim et al., 2018b). All of the significant resuscitation results found from the agar plates were confirmed by single-cell experiments. In addition, all of the strains used in this work were tested and shown to produce roughly the same number of persister cells (so there is no defect in the formation of persister cells) and shown to grow similar to the wild-type (so there is no growth defect).

### Persister cells wake by recognizing Ala

To elucidate insights about waking using defined medium, we hypothesized that a single amino acid could wake *E. coli* persister cells since *L*-alanine revives spores of *Bacillus subtilis* (Mutlu et al., 2018). Initially, four groups of five amino acids were tested (#1: Ala, Arg, Cys, Phe, Ser; #2: Gly, His, Thr, Val, Tyr; #3: Asn, Ile, Lys, Pro, Trp; and #4: Asp, Glu, Gln, Leu, Met) by plating *E. coli* persister cells on M9 agar plates with the amino acid combinations as the sole carbon source and checking for growth after four days (**Supplemental Fig. 1**). Larger colonies were obtained from group #1 compared to groups #3 and #4, indicating faster waking, and persister cells did not wake for group #2 (**Supplemental Fig. 1**). As expected, there was no persister cell growth with M9 agar that lacks amino acids (negative control). For the two best groups (#1 and #3), each amino acid was tested separately for persister cell waking, and clearly substantially more persister cells were revived with Ala (**Supplemental Fig. 2**).

To confirm that the macroscopic colonies seen on M9 agar plates reflected faster waking, single *E. coli* persister cells were observed on M9 5X Ala agarose gel pads using light microscopy. We found that 18 ± 1% of persister cells began to wake, as evidenced by dividing or elongating within 6 hours (**Fig. 1**, statistics shown in **Table S1**). As a negative control, single cell waking was monitored using Asn (5X) from group #3. Unlike with Ala, only 2 ± 2 % divided after 6 hours (**Fig. 1**, **Table S1**). Hence, persister cells with Ala wake 9-times more frequently than those with Asn. As an additional negative control, waking on agarose gel pads that lack amino acids (and any other carbon source) was investigated, and we found no persister cells wake after 6 h (**Fig. 1**). This lack of persister cell waking without nutrients also confirms that the persister cells used are *bona fide* dormant cells since, in contrast, exponential cells can wake in 6 h on agarose pads that lack nutrients (12 ± 4%, **Table S1**).

**Fig. 1.**
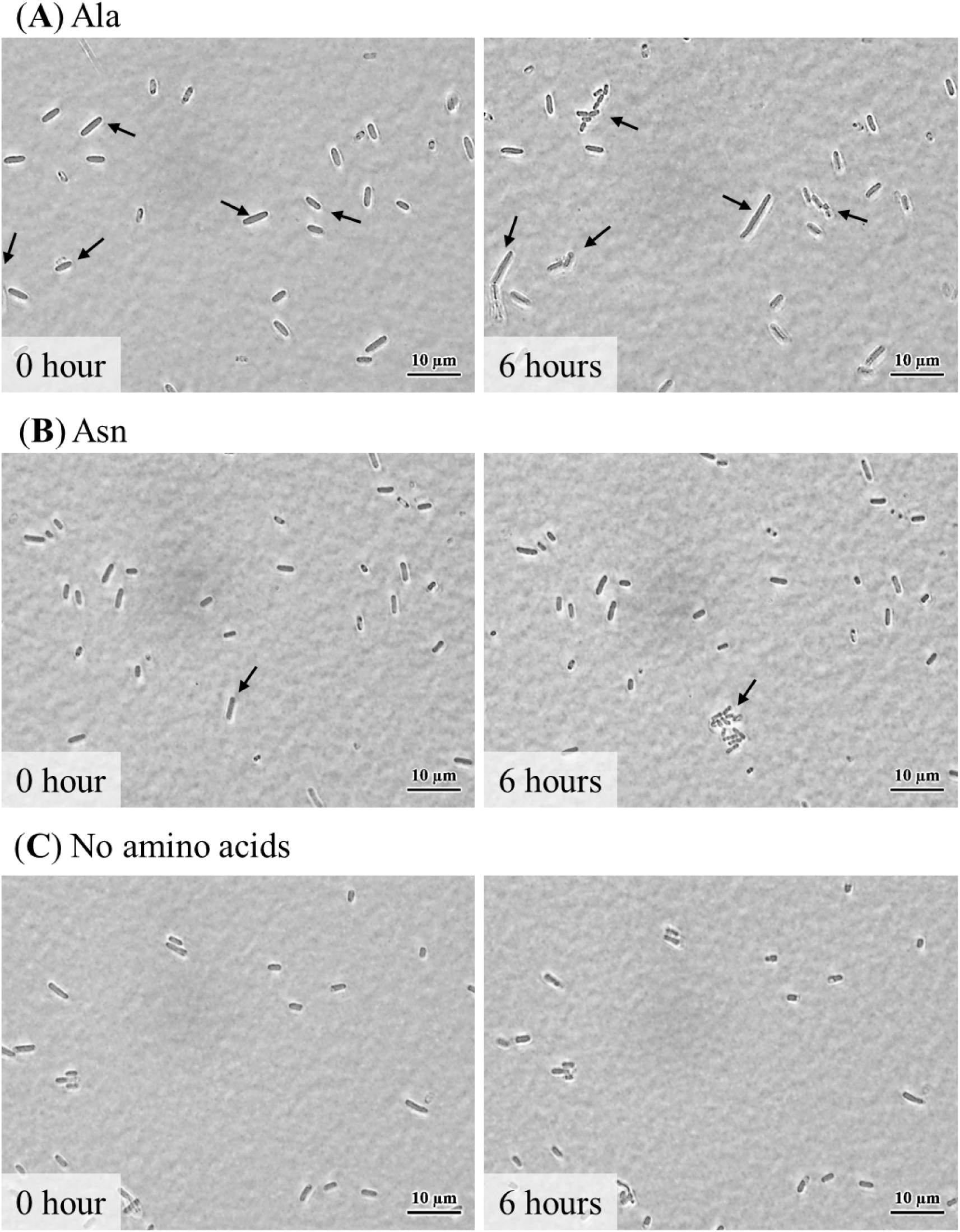
Single persister cells wake with Ala. *E. coli* BW25113 persister cells were incubated at 37 °C on M9 5X Ala agarose gel pads and images were captured at 0 h and 6 h. Black arrows indicate cells that resuscitate. Scale bar indicates 10 µm. Representative results from two independent cultures are shown. Tabulated cell numbers are shown in **Table S1**.

### Persister cells wake primarily by sensing nutrients rather than spontaneously

Along with waking as a response to a change in environmental conditions (e.g., the addition of nutrients (Kim et al., 2018b)), it has been reported that persister cells wake spontaneously (Balaban et al., 2004); hence, we investigated whether persister cells wake in the absence of Ala. To test this, single persister cells were observed on NaCl, NaCl + ampicillin, and NaCl + ampicillin + Ala agarose gel pads. The rationale is that if the persisters wake, ampicillin will lyse the cells (Uehara et al., 2009) which is seen as a disappearing single cell. We used NaCl buffer rather than M9 buffer so that the effect of ampicillin and Ala would be more clear.

We found that persister cells wake primarily in the presence of Ala since there was 4.4-fold more waking followed by lysis with Ala + ampicillin in 6 hours compared to gel pads that lacked Ala but contained ampicillin (**Fig. 2**, statistics in **Table S2**). As expected, for the negative control (NaCl only), there was no cell death seen in 6 hours (**Fig. 2A**). Therefore, most persister cells do not resuscitate spontaneously but instead sense the carbon source through some pathway. Furthermore, these results suggest the cells do not determine whether the antibiotic is present or not.

**Fig. 2.**
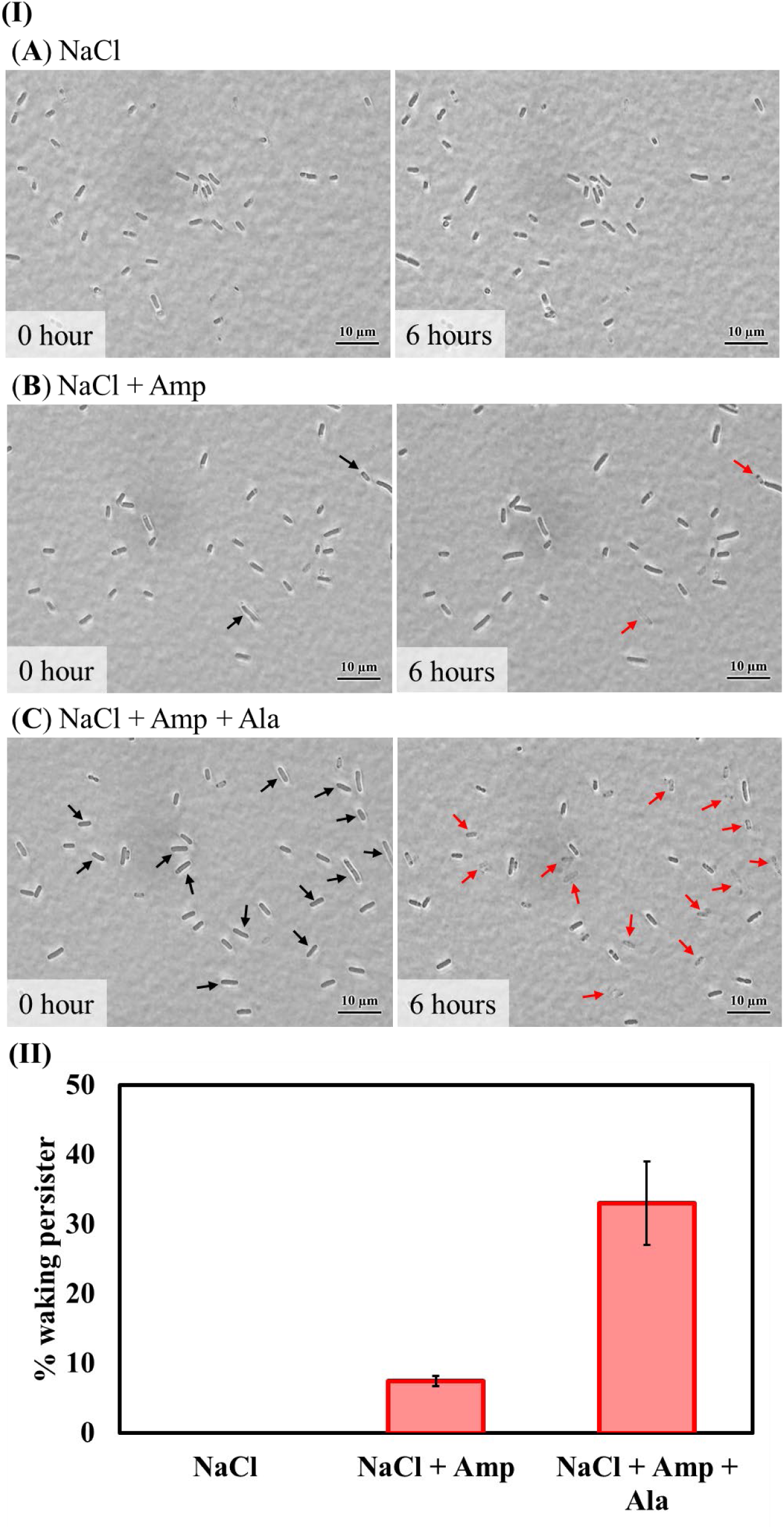
Single persister cells wake primarily by sensing nutrients rather than spontaneously. *E. coli* BW25113 persister cells were incubated at 37 °C on NaCl, NaCl + ampicillin (Amp), and NaCl + Amp + Ala agarose gel pads for 6 hours. **Panel I**: Cells first resuscitate (black arrows) and then lyse in the presence of ampicillin (red arrows). Scale bar indicates 10 µm. Representative results from two independent cultures are shown. **Panel II:** Persister cell waking (%) after 6 hours incubation. Tabulated cell numbers are shown in **Table S2**.

### Alanine is a waking signal

Since persister cells wake in the presence of Ala, we investigated whether Ala acts as a true signal rather than as a carbon and energy source by utilizing an isogenic *dadA* mutation that abolishes growth on alanine by inactivating *D*-amino acid dehydrogenase (Franklin and Venables, 1976; Wild and Klopotowski, 1981). We confirmed the *dadA* mutation abolishes growth on Ala (**Supplemental Fig. 3**) and found 4% of the single *dadA* cells wake with Ala whereas no cells wake without Ala (**Supplemental Fig. 3**, statistics in **Table S3**); hence, Ala wakes persister cells in the absence of growth. Moreover, we tested whether Ala wakes persister cells in the presence of the additional carbon and energy source pyruvate (final conc. 0.24 wt%) when Ala is not used for growth (i.e., with the *dadA* mutant). We found pyruvate does not wake single *dadA* persister cells whereas Ala wakes *dadA* persister cells, in the presence of pyruvate, to the same extent as in the absence of pyruvate (**Supplemental Fig. 3**, **Table S3**). Therefore, Ala is true signal for waking persister cells.

**Fig. 3.**
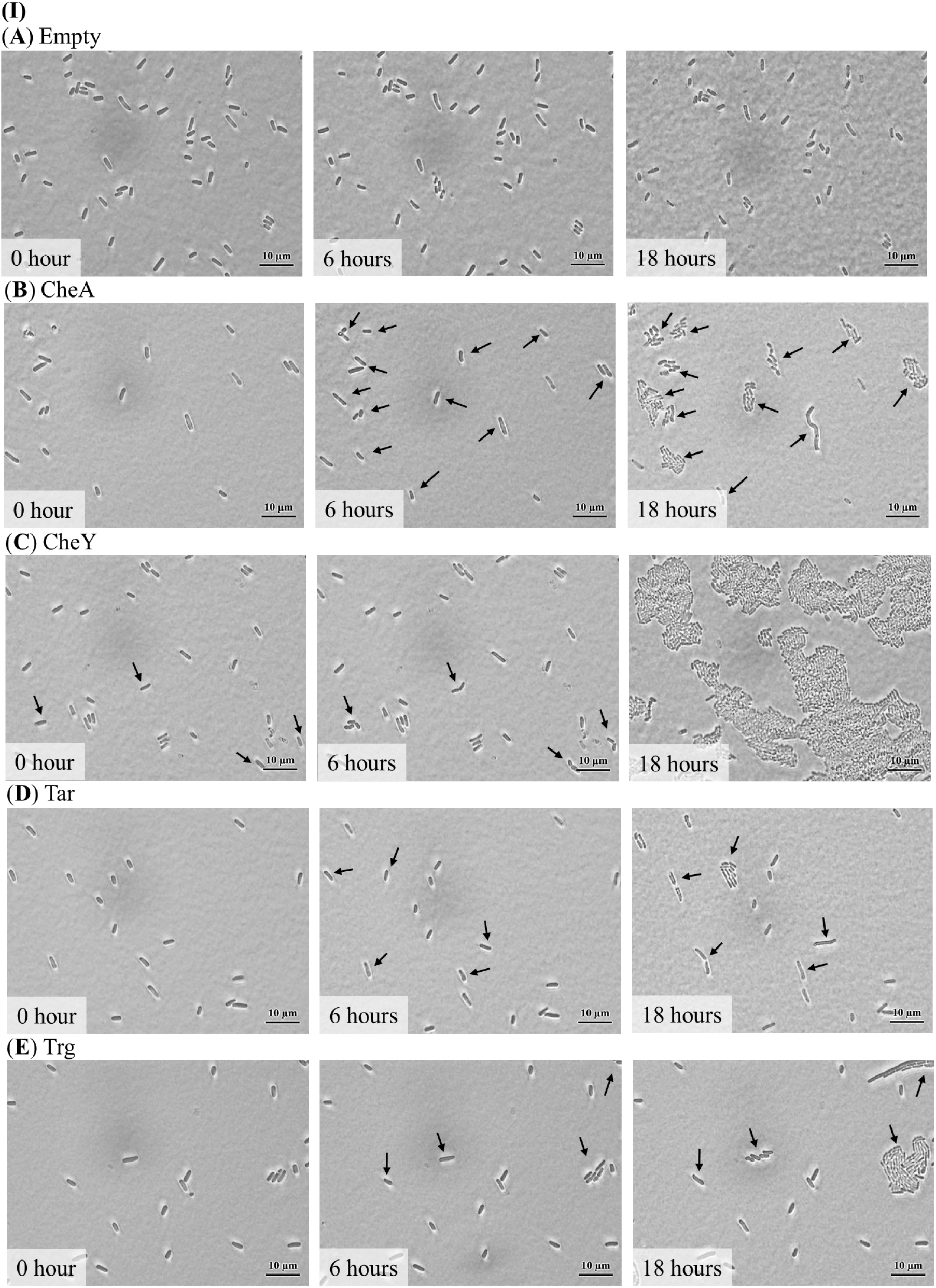
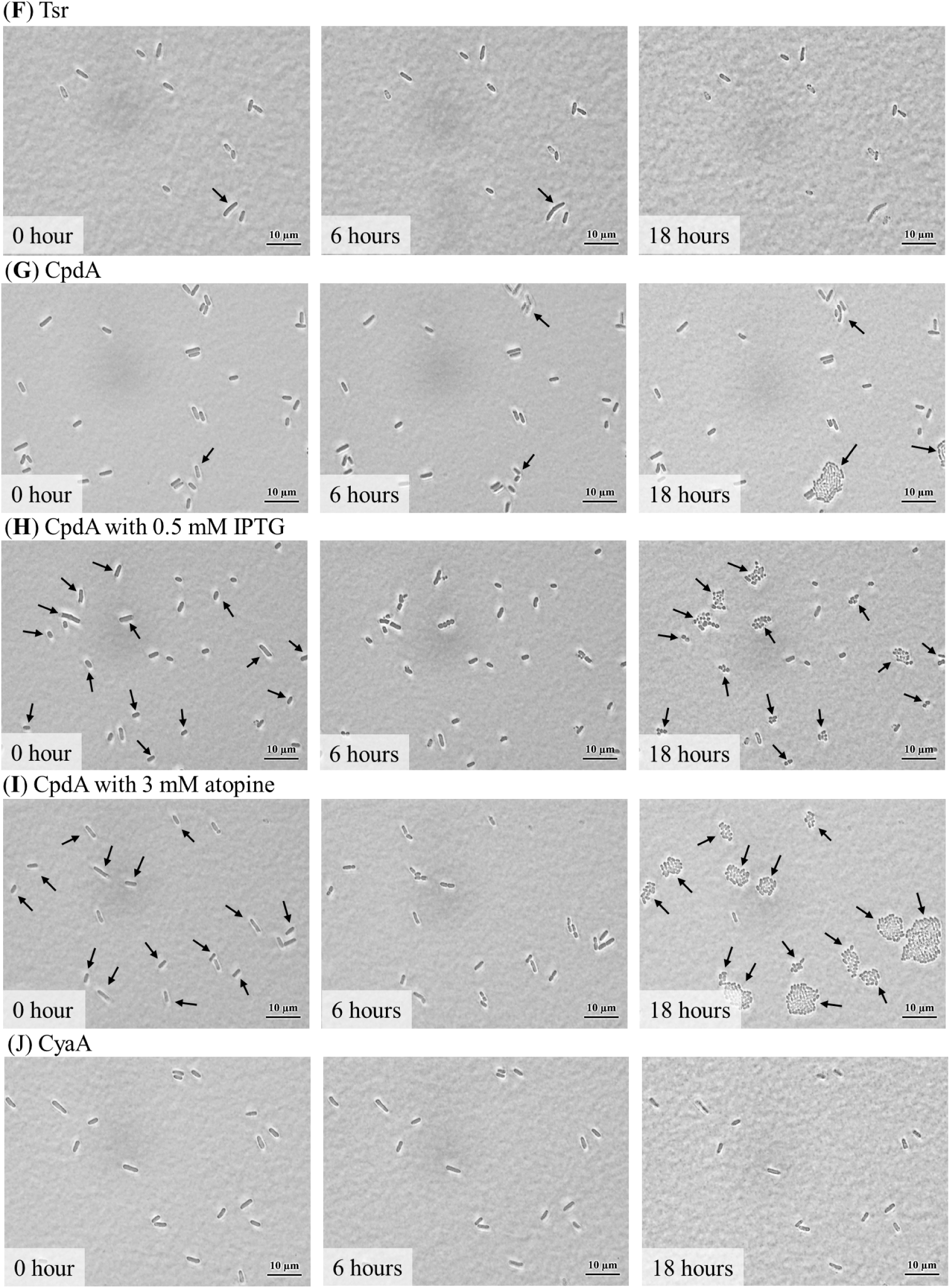
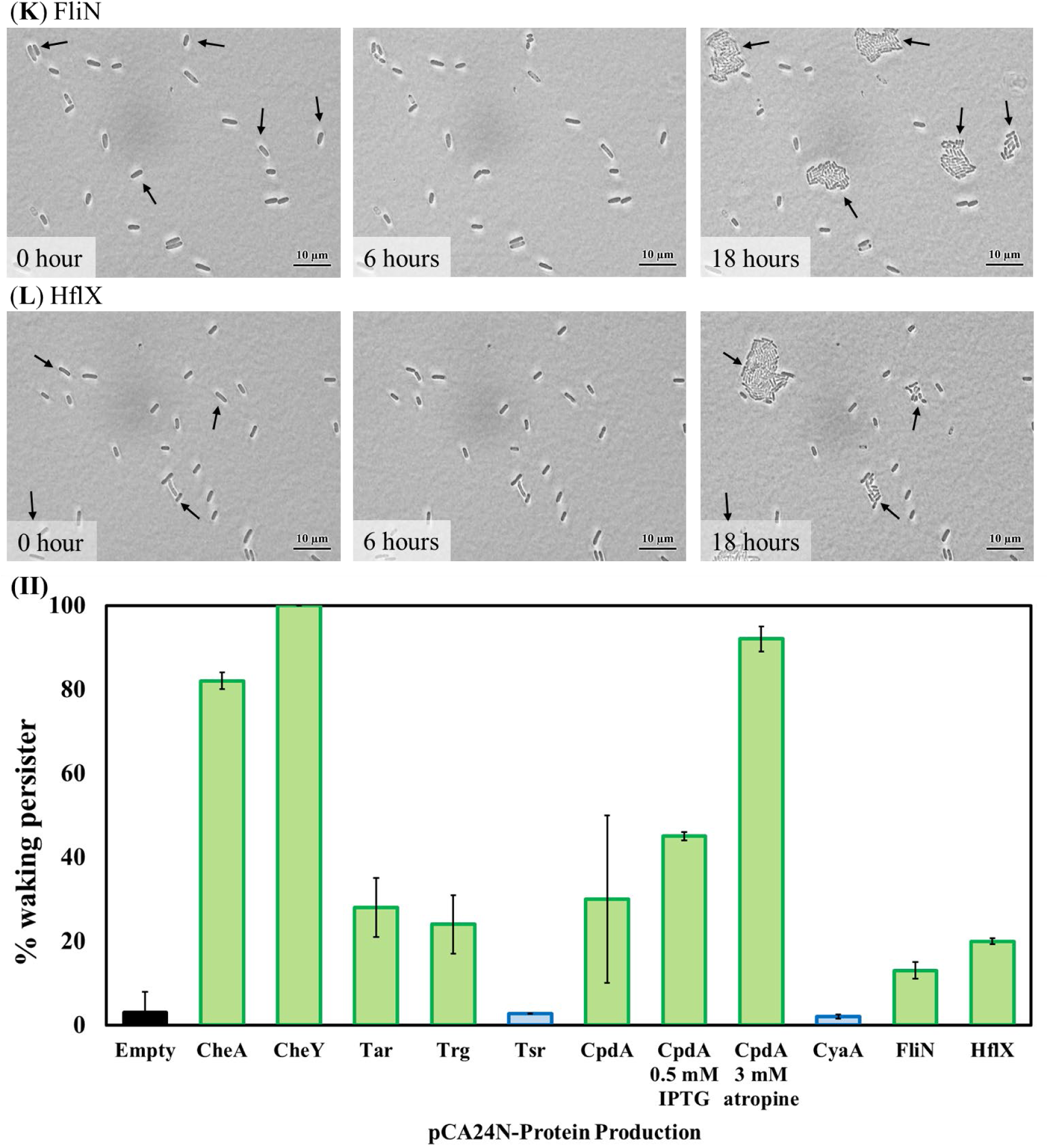
**Single persister cell waking on Ala after producing chemotaxis-related, cAMP-related, and ribosome-resuscitation-related proteins. Panel I**: *E. coli* BW25113 persister cells containing pCA24N, (**B**) pCA24N_*cheA*, (**C**) pCA24N_*cheY*, (**D**) pCA24N_*tar*, (**E**) pCA24N_*trg*, (**F**) pCA24N_*tsr*, (**G**) pCA24N_*cpdA*, (**H**) pCA24N_*cpdA* induced by 0.5 mM IPTG, (**I**) pCA24N_*cpdA* with 3 mM atropine, (**J**) pCA24N_*cyaA*, (**K**) pCA24N_*fliN*, and (**L**) pCA24N_*hflX* were incubated at 37 °C on M9 5X Ala agarose gel pads for 18 h. Black arrows indicate cells that resuscitate. Scale bar indicates 10 µm. Representative results from two independent cultures are shown. Prior to forming persister cells, cultures were grown without IPTG with 30 µg/mL Cm. **Panel II**: Persister cell waking (%) after incubating for 18 h. Tabulated cell numbers are shown in **Table S4**.

### Persister cells resuscitate through the chemotaxis apparatus

Since most persister cells are waking by responding to the presence of Ala as a signal, we investigated the mechanism of how persister cells resuscitate by looking for faster waking when individual proteins are produced. Hence, we pooled all the ASKA plasmids (each pCA24N-derived plasmid produces one *E. coli* protein) and electroporated them into *E. coli* BW25113, formed persister cells, and selected for faster growth on M9 5X Ala agar plates containing chloramphenicol to retain the plasmid. After 3 days, some colonies could be seen and after 8 days, different phenotypes were clearly seen (**Supplemental Fig. 4**). The faster (bigger) colonies were purified and sequenced and six proteins were identified: PsiF, PanD, YmgF, YjcF, PptA, and CheY.

For these six putative proteins that stimulate persister cell waking, single cells harboring the ASKA plasmid were observed on M9 5X Ala agarose gel pads and compared with persister cells with an empty plasmid (*E. coli* BW25113/pCA24N) (**Supplemental Fig. 5, Table S4**). Note that due to the metabolic burden of the plasmid, the cells wake slower so the microscopic observations were extended to 18 h. We found single cells that produce PanD (23-fold) and YmgF (18-fold) wake at a higher frequency with Ala, compared to cells containing the empty plasmid (**Table S4**). Remarkably, all the single persister cells that produced the chemotaxis response regulator CheY resuscitated (33-fold enhancement, **Fig. 3, Table S4**), and 82% of the single persister cells that produced response regulator CheA resuscitated (27-fold enhancement) (**Fig. 3, Table S5**); In addition, single cells of deletion mutants Δ*cheY* and Δ*cheA* are completely inhibited in their waking on M9 Ala (**Supplemental Fig. 6**). Critically, the CheY and CheA strains do not have defects in either persister cell formation and growth (**Table S6**). Together, these results demonstrate the importance of the chemotaxis system in nutrient recognition and waking in persister cells.

To investigate further the link between chemotaxis and persister cell waking, we tested all five of the methyl-accepting chemotaxis proteins (Tsr, Tar, Trg, Tap, and Aer) for waking on agar plates by using isogenic mutants and found inactivating Tar and Trg decreased waking compared to the wild type (**Supplemental Fig. 7**). Corroborating these results, persister cells that produce Tar and Trg woke faster than those with the empty plasmid and faster than cells producing Aer, Tsr, and Tap (**Supplemental Fig. 8**).

Single cells producing Tar (9-fold) and Trg (8-fold) also woke with a higher frequency than cells containing the empty plasmid, and single cells producing Tsr woke the same as those with the empty plasmid (**Fig. 3, Table S5**). Hence, persister cells wake on alanine via the chemotaxis system utilizing methyl-accepting chemotaxis proteins Tar and Trg and response regulators CheY and CheA.

### Persister cell waking does not require flagellar rotation

Since the chemotaxis system involves the flagellum, we investigated its role in persister cell waking. Using agar plates, there was no effect of the flagellum as tested by inactivating the flagella motor MotA (**Supplemental Fig. 9**), and single persister cells with inactivated flagella motor MotB did not wake significantly differently than wild-type persisters (**Supplemental Fig. 10, Table S7**). Corroborating these results, inactivating the hook protein FlgE (Brown et al., 2012) had no effect on persister waking (**Supplemental Fig. 11**), so flagellum movement is not required for persister cells waking. Note the flagella system has been linked previously to persister cell formation (Shan et al., 2015) but not to resuscitation.

In contrast, inactivating FhlD and FliN reduced the frequency of single cell waking by 3-fold (**Supplemental Fig. 10, Table S7**). In addition, producing FliN in single cells increased the waking frequency by 4.3-fold (**Fig. 3, Table S5**). Hence, the master transcriptional regulator of flagellar proteins is involved in persister cell waking.

### Persister cells wake by lowering cAMP levels

Having determined that persister cells wake by utilizing the chemotaxis apparatus, we explored how the external nutrient signal was propagated beyond the chemotaxis and flagellum system. Since cells can sense their environment (e.g., surfaces) by cAMP (Lee et al., 2018), and cAMP has been linked to persistence (Kwan et al., 2015b), we reasoned that persister cell waking may involve this secondary message.

To test the importance of cAMP, we produced the only adenylate cyclase in *E. coli*, CyaA (Tuckerman et al., 2009) via BW25113/pCA24N-*cyaA* and found using agar plates that cells with elevated cAMP have reduced waking compared to cells with the empty plasmid (**Supplemental Fig. 12**). Consistently, cells producing the cAMP-specific phosphodiesterase CpdA(Imamura et al., 1996) via BW25113/pCA24N-*cpdA* have increased waking on M9-Ala agar plates; we showed previously CpdA can eliminate cAMP by reducing its concentration 323-fold(Kwan et al., 2015b). Corroborating these results, cells lacking CpdA (BW25113 Δ*cpdA*) have reduced waking on M9-Ala plates (**Supplemental Fig. 12**).

Verifying these agar plate results, single cells producing adenylate phosphodiesterase CpdA before persister formation had a 15-fold greater (0.5 mM IPTG) and 8-fold greater (0 mM IPTG) frequency of waking (**Fig. 3, Table S5**) by reducing cAMP levels. In addition, producing adenylate cyclase CyaA reduced the frequency of waking by 30% compared to the empty plasmid (**Fig. 3, Table S5**). In a consistent manner, increasing cAMP levels by inactivating CpdA completely eliminated single cell waking (**Supplemental Fig. 13**, Table S8) and decreasing cAMP levels by inactivating CyaA led to 4.3-fold greater frequency of waking (78%) (**Supplemental Fig. 13, Table S8**).

As additional proof of the impact of cAMP, we used the cAMP inhibitor atropine (Huynh et al., 2012) and found single cells wake with atropine (3 mM) at twice the frequency (**Supplemental Fig. 14**). Consistently, adding exogenous cAMP (2 mM) reduced the frequency of single-cell waking 3-fold (**Supplemental Fig. 14**). Moreover, combining atropine with production of CpdA to reduce cAMP increased the frequency of waking 4-fold such that 92% of the cells woke in 6 h (**Fig. 3, Table S5**). Therefore, reducing cAMP levels increases persister cell waking dramatically.

### Persister cells wake by resuscitating and rescuing ribosomes

Since reducing levels of cAMP led to faster resuscitation, we explored further the mechanism for their waking. We hypothesized that since elevated cAMP levels increase ribosome hibernation during starvation (Shimada et al., 2013), lowering cAMP levels should lead to ribosome resuscitation; hence, we tested the impact of changing the concentration of ribosome resuscitation factor HflX (Gohara and Yap, 2018); HflX also rescues ribosomes from mRNA (Zhang et al., 2015). We utilized ribosome rescue factor ArfB (Abo and Chadani, 2014) as a negative control since it is positively regulated by cAMP (Raghavan et al., 2011). In addition, since trans-translation is used to rescue ribosomes from mRNA that lack a stop codon (Abo and Chadani, 2014), we also investigated the impact of SsrA on persister cell waking, using the strain we constructed (Wang et al., 2009).

For M9 Ala agar plates, we found cells producing HflX woke faster, compared to cells with the empty plasmid whereas cells producing ArfB did not wake, indicating ArfB reduces waking dramatically (**Supplemental Fig. 15**). Similarly, for single cells, we found inactivating HflX completely inhibited waking (**Supplemental Fig. 16, Table S9**). In contrast, inactivating ArfB in single cells had no effect (**Supplemental Fig. 16, Table S9**). Corroborating these results, producing HflX in single cells increased the frequency of waking dramatically (7-fold) (**Fig. 3, Table S5**).

For ribosome rescue, using agar plates, we found inactivating SsrA led to significantly slower waking (**Supplemental Fig. 15**). Similarly, for single cells, inactivating SsrA almost completely inhibited waking (frequency of waking reduced to less than 1%) (**Supplemental Fig. 16, Table S9**). Therefore, resuscitating ribosomes via HflX and SsrA is vital for persister cell resuscitation. Previously, SsrA has been shown to be have a modest role (2- to 8-fold) in the formation of persister cells (Shi et al., 2013).

### Persister cells wake by recognizing sugars derived from biofilms

Using alanine to wake cells, it was not clear how cAMP levels are reduced; hence, we studied resuscitation with the building blocks of the biofilm matrix since we hypothesized that those materials would be readily available. Initially, we chose four polysaccharide building blocks of the most prevalent polymers of *E. coli* biofilms: *D*-glucose from cellulose (Klemm et al., 2005), *D*-glucosamine and *N*-acetyl-D-glucosamine from poly-*N*-acetylglucosamine (Pokhis et al., 2015), and α-*D-*glucose 1-phosphate from colonic acid (Patel et al., 2012; Stevenson et al., 1996). The final concentration of each sugar was kept at 0.4%, and each polysaccharide building block was utilized as the sole carbon and energy source.

Using minimal agar plates that contain these polysaccharides at 0.4 wt% and Ala at 0.04 wt% (0.4 wt% Ala inhibited waking,), we found that persister cells wake more rapidly on biofilm-derived polymers than on Ala since colonies form two days faster on plates with the polysaccharide precursors (**Supplementary Fig 17**). Since waking with glucose was most rapid, we focused on using it for persister resuscitation. As expected, we found the frequency of single cell waking was higher with glucose compared to Ala; for example, in 6 h, 92 ± 2% cells resuscitate on glucose compared to 18 ± 1% on Ala (**Fig. 4, Table S10**).

**Fig. 4.**
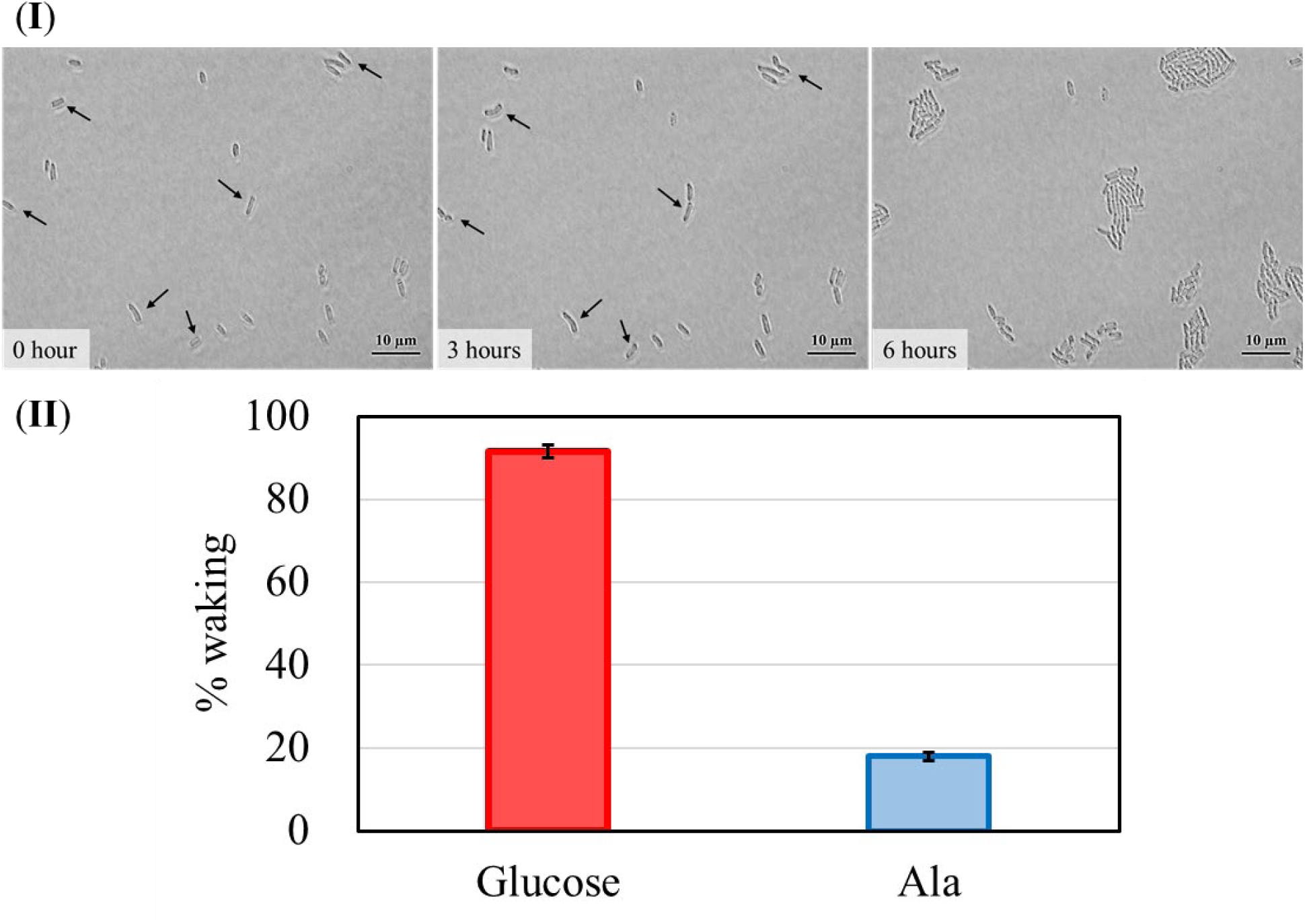
Single persister cells wake with glucose. **Panel I**: *E. coli* BW25113 persister cells were incubated at 37 °C on M9 0.4% glucose agarose gel pads and images were captured at 0, 3, and 6 h. Black arrows indicate cells that resuscitate. Scale bar indicates 10 µm. Representative results from two independent cultures are shown. Tabulated cell numbers are shown in **Table S1**. **Panel II**: Comparison of single *E. coli* BW25113 persister cells waking on minimal medium with 0.4% glucose and 0.04% alanine for 6 h.

### Glucose waking requires transporters PtsG and MglB and chemotaxis

For waking on Ala, we discovered that the methyl-accepting chemotaxis proteins Trg and Tar are used for persister resuscitation. Hence, we hypothesized that the glucose phosphotransferase transport protein PtsG, which interacts indirectly with the chemotaxis system via chemoeffector CheA (Neumann et al., 2012), may be responsible for waking with glucose. Supporting this hypothesis, we found that inactivating PtsG reduced waking on agar plates (**Supplementary Fig. 18**) and nearly abolished the frequency of single cell waking on glucose (70-fold reduction, **Fig. 5, Table S11**). In addition, production of PtsG from pCA24N-*ptsG* abolished glucose waking indicating the stoichiometry of the PtsG proteins in the inner membrane are critical (**Fig. 6, Table S12**). As a negative control, the frequency of waking was tested for the maltose PTS transporter by inactivating MalE; as expected, deletion of *malE* had no impact on waking with glucose (**Fig. 5, Table S11**). Hence, persister cell waking glucose transport through the phosphotransferase system.

**Fig. 5.**
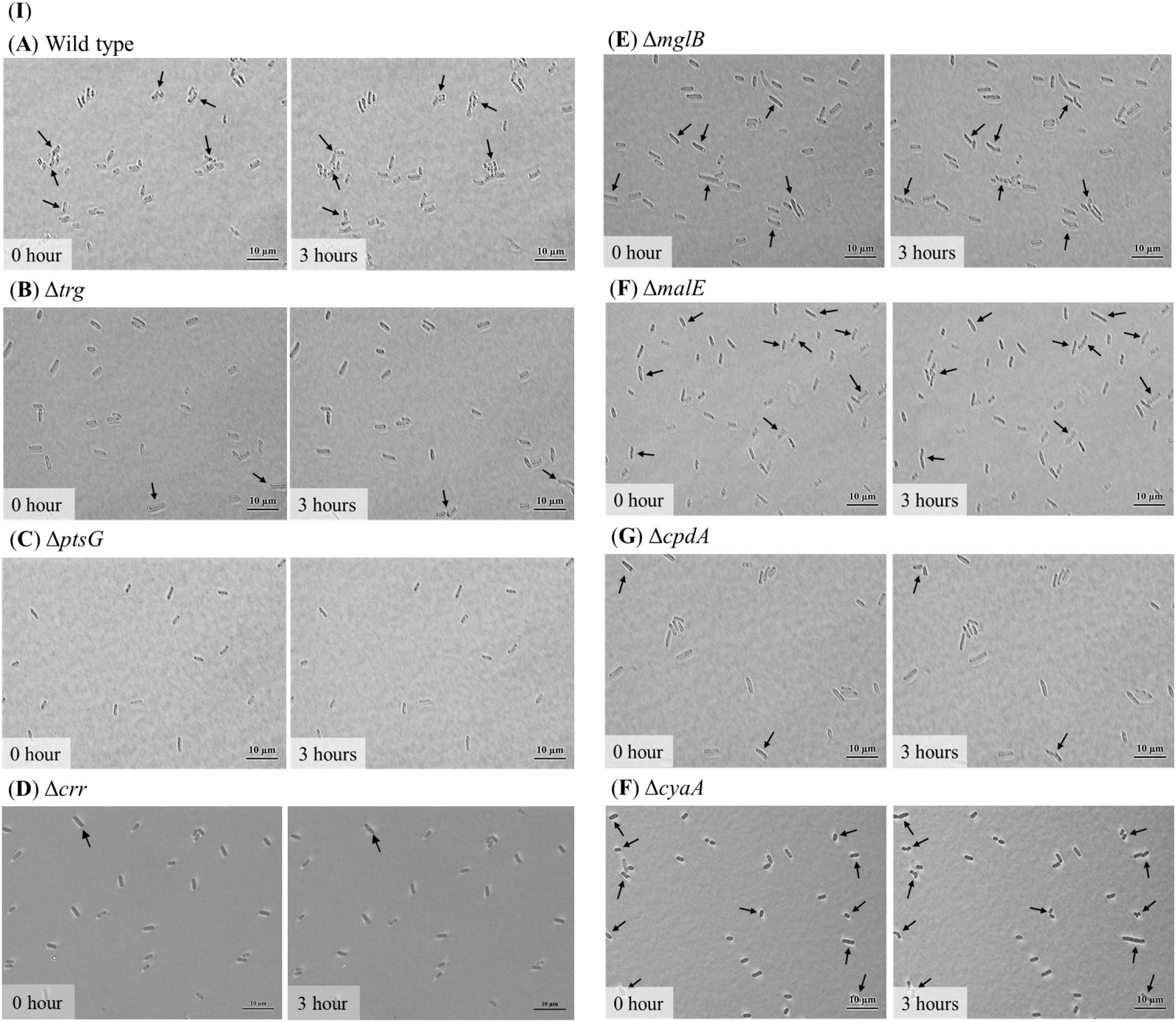
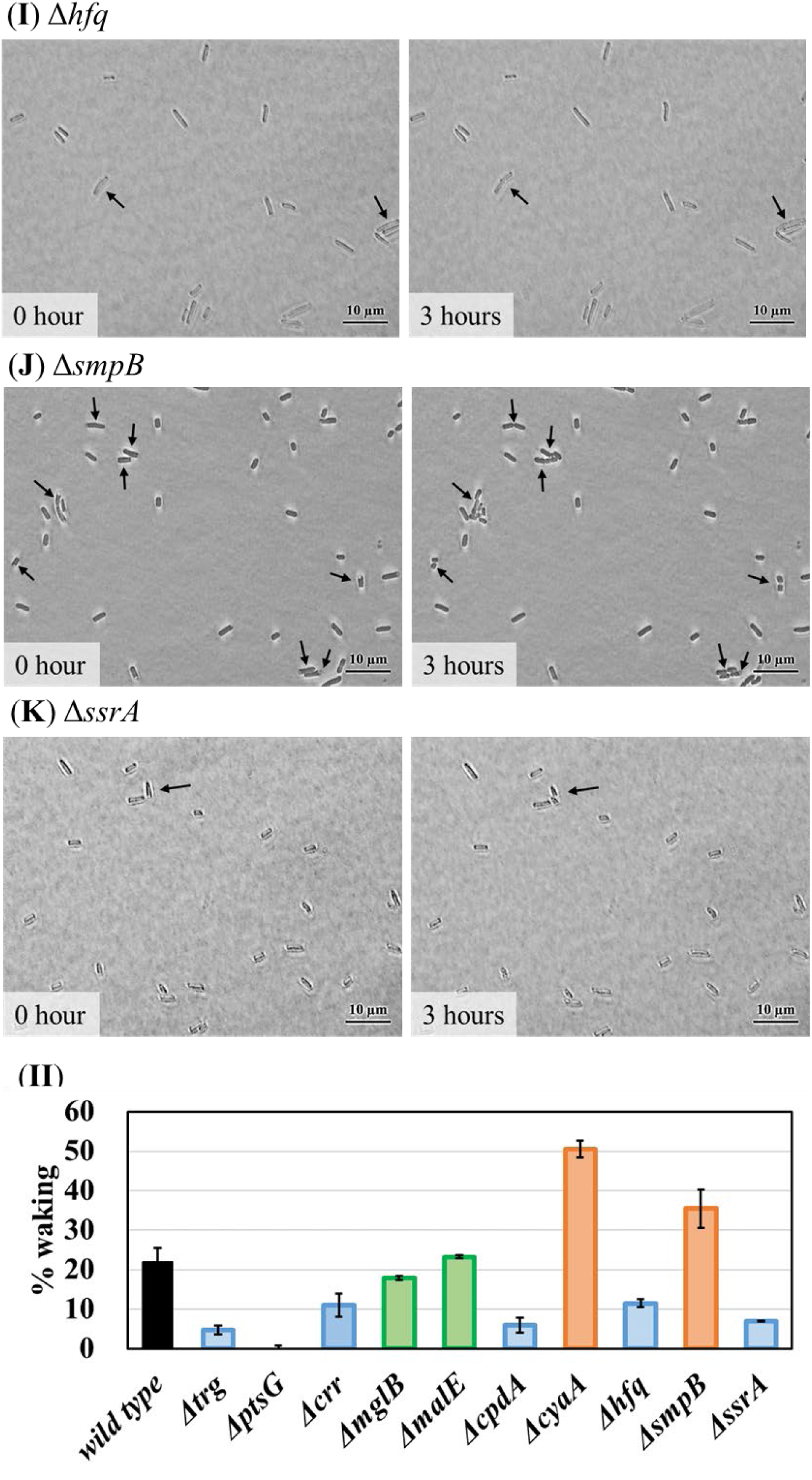
Single persister cell waking on glucose after inactivating proteins related to transport, cAMP, sRNA, and ribosome rescue. **Panel I**: *E. coli* BW25113 persister cells (**A**) and isogenic deletion mutants (**B**) Δ*trg*, (**C**) Δ*ptsG*, (**D**) Δ*crr*, (**E**) Δ*mglB*, (**F**) Δ*malE*, (**G**) Δ*cpdA*, (**F**) Δ*cyaA*, (**I**) Δ*hfq*, (**J**) Δ*smpB*, and (**K**) Δ*ssrA* were incubated at 37 °C on 0.4% glucose agarose gel pads for 3 h. Black arrows indicate cells that resuscitate. Scale bar indicates 10 µm. Representative results from two independent cultures are shown. **Panel II**: Persister cell waking (%) after incubating for 3 h. Tabulated cell numbers are shown in **Table S11**.

**Fig. 6.**
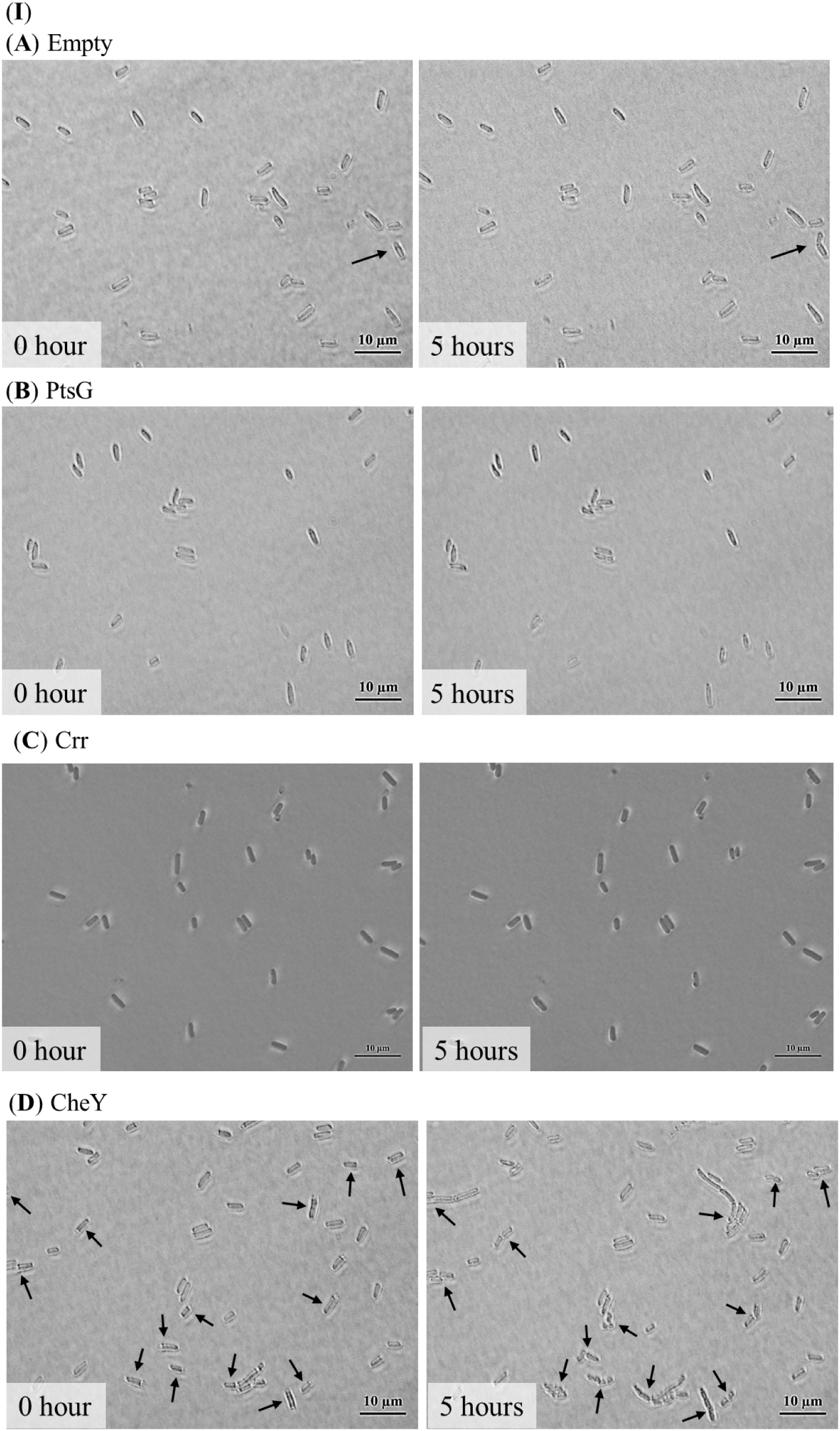
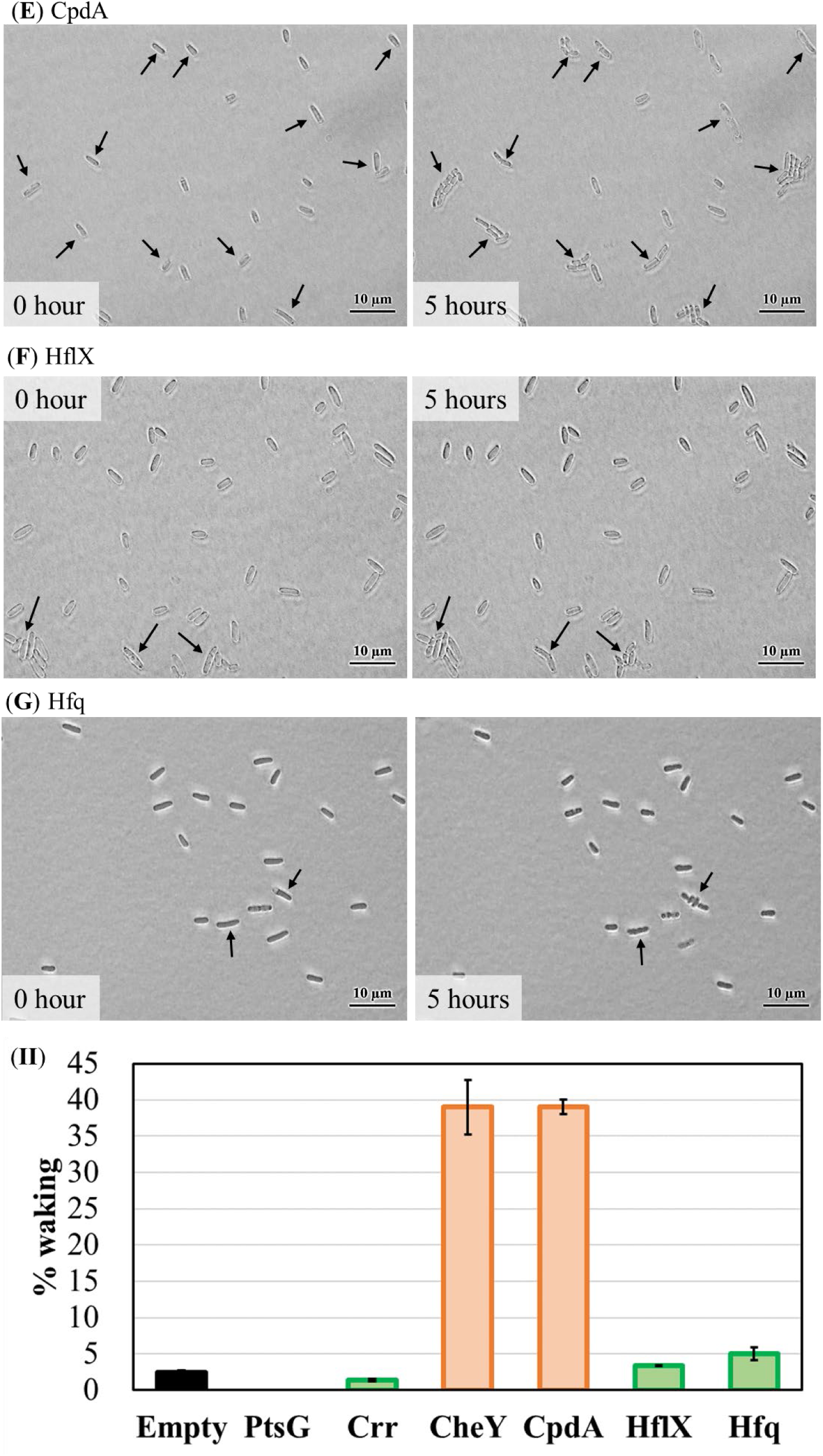
Single persister cell waking on glucose after producing proteins related to transport, chemotaxis, cAMP, ribosome resuscitation, and sRNA. **Panel I**: *E. coli* BW25113 persister cells containing (**A**) pCA24N, (**B**) pCA24N_*ptsG*, (**C**) pCA24N_*crr*, (**D**) pCA24N_*cheY*, (E) pCA24N_*cpdA*, (**F**) pCA24N_*hflX*, and (**G**) pCA24N_*hfq* were incubated at 37 °C on M9 0.4% glucose agarose gel pads for 5 h. Black arrows indicate cells that resuscitate. Scale bar indicates 10 µm. Representative results from two independent cultures are shown. Prior to forming persister cells, cultures were grown without IPTG with 30 µg/mL Cm. **Panel II**: Persister cell waking (%) after incubating for 5 h. Tabulated cell numbers are shown in **Table S12**.

Since persister waking on glucose occurs via PtsG, we tested whether the chemotaxis system was involved since the transported glucose stimulates chemotaxis (Neumann et al., 2012). By producing chemoeffector CheY, we found the frequency of single-cell waking increased dramatically (16-fold) (**Fig. 6, Table S12**). Hence, persister cell waking with glucose also depends on an active chemotaxis system.

We also tested whether the resuscitation of persister cells on glucose depends on the chemotaxis ABC glucose transporter MglB found a 1.2-fold reduction in the frequency of single-cell waking upon inactivating MglG (**Fig. 5**). Corroborating this result, deleting the methyl-accepting chemotaxis protein Trg, which interacts with glucose from MglB (Neumann et al., 2012), reduced the frequency of single-cell waking by 5-fold (**Fig. 5, Table S11**). Hence, persister cell resuscitation on glucose requires both well-known glucose transport systems as well as the chemotaxis system.

### Glucose waking requires reduced cAMP

Since Ala resuscitation requires the cell to lower cAMP levels, we explored if this occurs with glucose and found again that low cAMP levels are needed since adding 2 mM cAMP as well as inactivating phosphodiesterase CpdA (Imamura et al., 1996) to increase cAMP levels, reduces waking on glucose on agar plates (**Supplementary Fig. 19**). Corroborating these plate results, producing CpdA to lower cAMP levels increased the frequency of waking of single cells dramatically (16-fold) on glucose (**Fig. 6, Table S12**), and inactivating adenylate cyclase CyaA (Tuckerman et al., 2009) to eliminate cAMP, increased the frequency of waking on glucose substantially (2.3-fold) (**Fig. 5, Table S11**). In addition, inactivating CpdA reduced the frequency of single cell waking by 4-fold (**Fig. 5, Table S11**). Therefore, cAMP levels must be low for persister cells to wake with glucose.

### Glucose transport lowers cAMP concentrations

We hypothesized that the requisite reduction in cAMP upon resuscitation with glucose is a result of glucose transport with PtsG, since the associated phosphotransferase system enzyme EIIA^Glc^ becomes unphosphorylated; unphosphorylated EIIA^Glc^ inactivates adenylate cyclase CyaA and leads to low cAMP (Deutscher, 2008; Yao et al., 2011) (a well-known aspect of catabolite repression). In support of this model, we tested the *crr* isogenic mutant (*crr* encodes EIIA^Glc^) with single cells and found that inactivating EIIA^Glc^ reduced the frequency of single-cell waking 2-fold (**Fig. 5**). In addition, single cells producing EIIA (pCA24N-*crr*) had 0.6-fold reduced waking frequency (**Fig. 6**). Therefore, reducing cAMP concentrations through EIIA^Glc^ is important for persister cell waking.

### Glucose waking requires ribosome resuscitation

Since waking on Ala requires cells to rescue ribosomes from corrupt mRNA and from dimerization via HflX and to rescue ribosomes from corrupt mRNA via SsrA, we tested whether these proteins are also necessary for persister cell resuscitation on glucose. For waking on agar plates, inactivating trans-translation (i.e., SsrA) abolished waking (**Supplementary Fig. 20**). For single-cell waking with glucose, there was only a small positive effect of producing HflX (**Fig. 6, Table S12**); however, inactivating trans-translation (Δ*ssrA*) reduced waking dramatically (-3-fold) ((**Fig. 5, Table S11**). Therefore, ribosome resuscitation through SsrA is primarily responsible for waking with glucose.

### Cells wake and begin chemotaxis simultaneously

To determine if persister cells resuscitate and undergo chemotaxis simultaneously, we observed single persister cell waking inside motility agar with a glucose gradient; therefore, unlike the previous results where persister cells resuscitated on the surface of agarose gel pads, here persisters resuscitate and move planktonically inside a 0.3% agar (motility) gel. Note that the movement of the cells in these experiments is not due to fluid flow or random motion since the cell movement is always toward glucose in many repeated trials and since there is no fluid flow in this static 0.3% gel which is a semi-solid rather than a liquid.

Critically, we found that 20 ± 6% of the wild-type persister cells resuscitate and undergo chemotaxis immediately toward the glucose (**Supplementary video 1**). As expected, wild-type persister cells do not move without glucose. Since reducing cAMP by inactivating *cyaA* increases waking (**Fig. 5, Supplementary Fig. 13**), we hypothesized that the *cyaA* mutation would magnify this effect of instantaneous waking; strikingly, 94 ± 2% of the *cyaA* persister cells moved immediately toward glucose, indicating that the reduction of cAMP primes the cells for waking and chemotaxis. Furthermore, we reasoned that inactivating EIIA would also reduce cAMP and stimulate waking and found 96 ± 6% of the *crr* cells woke and began chemotaxis to glucose (**Supplementary video 1**). Critically the *crr* mutant cells wake instantaneously without glucose and show random movement in the M9 motility agar that lacks glucose confirming that low cAMP levels lead to more active resuscitation. The *crr* mutants still formed persister cells since any non-persister cells were removed during the 3 h ampicillin treatment. Therefore, the persister cells that resuscitate immediately begin moving toward nutrients.

## DISCUSSION

Spores are the ultimate bacterial resting state, and *L*-alanine revives spores of *Bacillus subtilis* (Mutlu et al., 2018). Hence, we hypothesized that *L*-alanine or another amino acid may wake *E. coli* persisters. By investigating all 20 amino acids, we found that *L*-alanine wakes *E. coli* persister cells much faster than other amino acids (**Supplemental Fig. S1, S2**). To determine insights into how *L*-alanine wakes *E. coli* persisters, we pooled the ASKA clones, isolated their plasmids, electroporated them into *E. coli* BW25113, made persister cells, and selected for faster waking. From this selection, six proteins were identified that increased waking: PsiF, PanD, YmgF, YjcF, PptA, and CheY. From these six proteins, we found PanD and YmgF increase waking 20-fold. PanD (aspartate 1-decarboxylase) (Cronan, 1980) which catalyzes the reaction of *L*-aspartate to β-alanine; hence, PanD produces an alanine-like compound. YmgF is related to an inner membrane division septum protein (Karimova et al., 2009); therefore, the persister cells wake and are primed for cell division. Critically, we determined the chemotaxis response regulator CheY and the chemotaxis methyl-accepting chemotaxis proteins Tar and Trg are utilized for persister resuscitation. Surprisingly, the chemotaxis methyl-accepting protein Tsr, which is primarily responsible for Ala chemotaxis (Tajima et al., 2011), was not involved in persister waking; however, Tar is also utilized for Ala chemotaxis (Yang et al., 2015). Therefore, persister cells wake through the same nutrient sensors they utilize for chemotaxis. Once resuscitated, the cells then rapidly move toward nutrients since the chemotaxis system is primed immediately, and the lack of nutrients (and energy for protein production) is what caused the dormancy in the first place (Kwan et al., 2013).

cAMP activates ribosome hibernation as 100S dimers during starvation by activating ribosome-binding protein RMF (Shimada et al., 2013) so it makes sense for cells to reverse this process (as we find here) and reduce cAMP levels to wake in the presence of nutrients, since we identified that the activity of ribosomes dictates the speed of persister resuscitation (Kim et al., 2018b). Supporting this insight, we found that inactivation of the ribosome rescue factor HflX (heat shock-induced ribosome-dependent GTPase) completely stopped waking of single cells and production of HflX increased the frequency of waking dramatically; *hflX* (in the same operon as *hfq*) is repressed by cAMP-Crp (Lin et al., 2011). In contrast, ribosome rescue factor Arfb (peptidyl-tRNA hydrolase) had no effect because it is induced by cAMP-Crp (Raghavan et al., 2011). Critically, all bacteria have mechanisms for rescuing ribosomes since without rescue, all ribosomes would be depleted in less than one generation (Moore and Sauer, 2007), and we found that without ribosome rescue via the trans-translation system (SsrA), persister cell waking is almost completely inhibited for Ala (**Supplementary Fig. 16**) and reduced for glucose (**Fig. 5**). Hence, ribosome rescue during persister cell resuscitation may be a general mechanism and should provide ample ribosomes for resuscitation.

Although it is counter intuitive, in regard to the *formation* of persister cells, high concentrations of cAMP allow ribosomes to hibernate and prepare the cell for nutrient-depleted conditions; yet, high concentrations of cAMP actually *reduce* persistence (Kwan et al., 2015b) (reduce cell dormancy). This is because cells with high cAMP concentrations are more prepared for stress and have less need to become dormant; i.e., more fit cells have dramatically less persistence (Hong et al., 2012).

Overall, by studying persister cell resuscitation with alanine, we have identified that (i) persister cells wake primarily in response to nutrient stimulus (rather than spontaneously), (ii) Ala is the most suitable amino acid signal for waking *E. coli* persister cells, (iii) this nutrient signal is perceived through the chemotaxis system via CheY, Trg, and Tar, (iv) second messenger cAMP concentrations are reduced for waking, and (v) ribosomes are restored for waking (**Fig. 7**). By studying persister cell resuscitation with glucose, we confirmed that chemotaxis and ribosome rescue are important for persister waking. Critically, the phosphotransferase system that imports glucose serves to reduce cAMP concentrations by dephosphorylating EIIA^Glc^, which allows ribosomes to resuscitate. Notably, the same mechanism for cAMP reduction is possible for alanine resuscitation via the chemotaxis system which detects it, since CheA, which we found to be critical for resuscitation with alanine, interacts directly with unphosphorylated EIIA^Glc (^Neumann et al., 201^2^). Therefore, persister cell resuscitation is an elegant (i.e., highly-regulated) response in which nutrients are perceived as environmental signals via membrane receptors; this environmental signal is propagated to ribosomes via the secondary metabolite cAMP, and persister cells commence foraging for food as the wake.

**Fig. 7.**
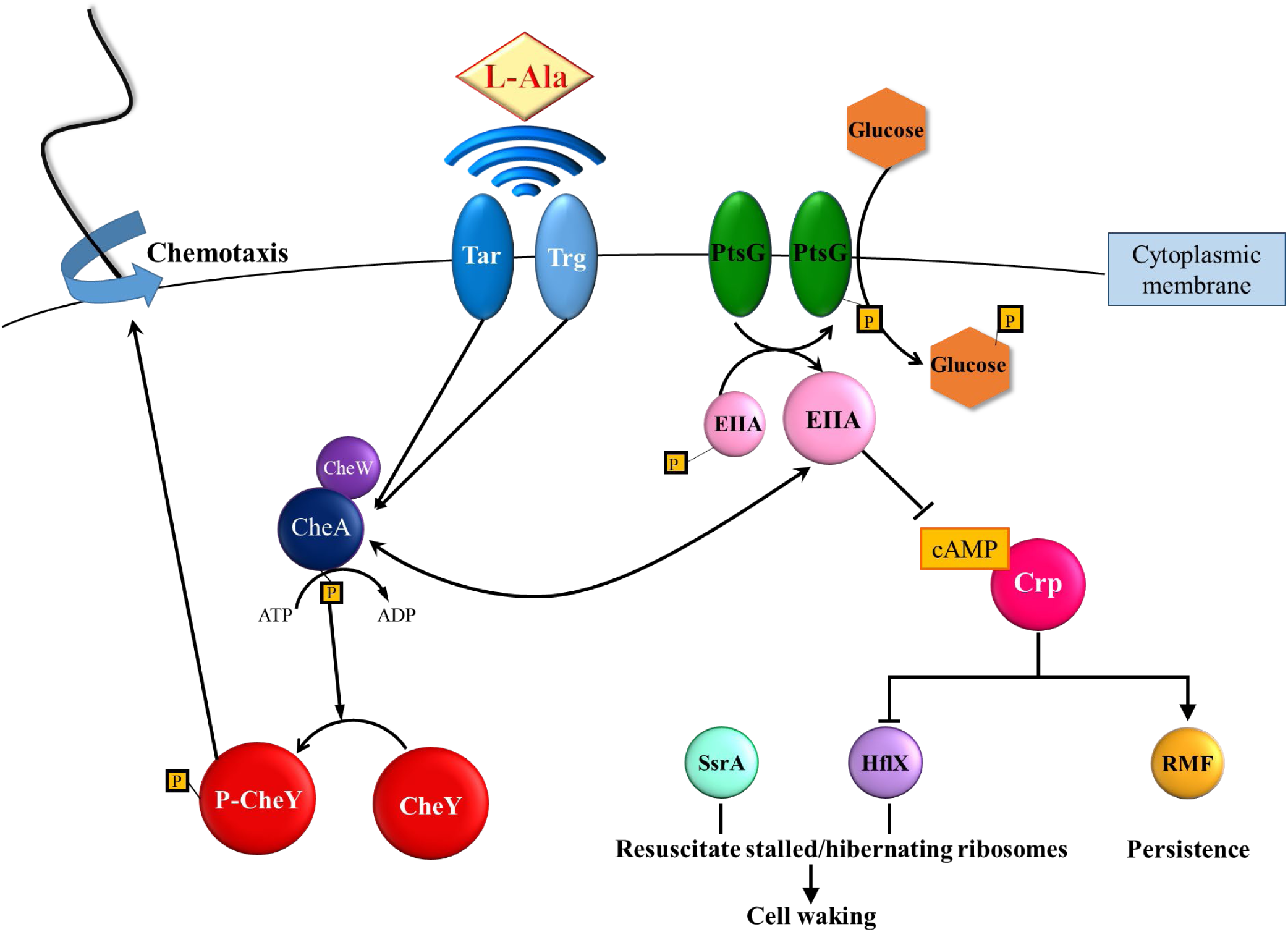
Schematic of persister cell waking via alanine and glucose. For alanine resuscitation, methyl-accepting chemotaxis proteins Tar and Trg sense the amino acid and relay this to chemotaxis response regulators CheA and CheY, which stimulate chemotaxis. For glucose resuscitation, phosphotransferase protein PtsG imports the sugar, which results in dephosphorylation of EIIA, reduction in cAMP, activation of chemotaxis, and ribosome rescue via HflX and SsrA. Spheres indicate proteins, diamonds indicate amino acids, hexagons indicate glucose, 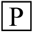 indicates phosphate, → indicates induction, and ⟞ indicates repression.

## METHODS

### Bacterial strain and growth conditions

The bacterial strains and plasmids used in this study are listed in **Table S13**. *E. coli* BW25113 and its isogenic mutants (Baba et al., 2006), and the pCA24N-based plasmids from the *E. coli* ASKA collection (Kitagawa et al., 2005) were used. Lysogeny broth (Sambrook et al., 1989) was used for routine cell growth, and M9 medium supplemented with amino acids (Rodriguez and Tait, 1983) was used for waking studies; strains were grown at 37 °C. Each of the 20 amino acids stock solutions were prepared as indicated in **Table S14** (Rodriguez and Tait, 1983).

### Observation of persister cells on agarose gel pads

We utilized our previous method for converting up 70% of the cell population into persister cells (Kim et al., 2018a; Kim et al., 2018b; Kwan et al., 2013): exponentially-growing cells (turbidity of 0.8 at 600 nm) were treated with rifampicin (100 µg/mL for 30 min) to form persister cells by stopping transcription and non-persister cells were removed by lysis via ampicillin treatment (100 µg/mL for 3 h). Cells were harvested at 17,000 g for 1 min and washed with 1x phosphate buffered saline buffer (Dulbecco and Vogt, 1954) (PBS) buffer twice to remove all possible carbon sources, then re-suspended with 1 mL of 1x PBS. Gel pads of 1.5% agarose were prepared (Kim et al., 2018b) and 5 µL of persister cells were added, kept at 37°C, and observed using a light microscope (Zeiss Axio Scope.A1, bl_ph channel at 1000 ms exposure) every hour.

### Observation of persister cells on motility gel pads

An M9-gradient glucose motility gel pad (0.3% agar) was prepared by solidifying for 1.5 hours and adding 10 μL of 5 wt% glucose and 10 μL of persister cells. The movement of the persister cell was observed using light microscopy (Zeiss Axio Scope.A1, bl_ph channel at 1000 ms exposure).

### Persister resuscitation screen

All 4,267 ASKA clones (GFP-) (Kitagawa et al., 2005) were combined, grown to a turbidity of 2 at 600 nm in LB medium, and their plasmids isolated using a plasmid DNA Mini Kit I (OMEGA Bio-tek, Norcross, GA, USA). The pooled ASKA plasmids (1 µL containing 30 ng of DNA) was electroporated into 50 µL of *E. coli* BW25113 competent cells, 1 mL LB medium was added, and the cells were grown to a turbidity of 0.5 in LB medium. Chloramphenicol was added (final conc. 30 µg/mL) to the culture, and the cells were incubated at 250 rpm to a turbidity of 0.8. Rifampicin followed by ampicillin was added to make persister cells, the persister cells were washed twice with 1x PBS buffer, diluted 100-fold 1x PBS buffer, and 100 µL was plated on M9 5X Ala (Cm) agar plates and incubated at 37°C; faster colony appearance indicated faster persister resuscitation.

## AUTHOR’S CONTRIBUTIONS

RY and SYS performed the experiments. RY, SYS, MJB, and TKW designed the experiments.

## AVAILABILITY OF DATA AND MATERIALS

All data are present in the manuscript.

## ACKNOWLEDGEMENTS

This work was supported by funds derived from the Biotechnology Endowed Professorship at the Pennsylvania State University. We thank Prof. John Parkinson and Prof. Pushkar Lele and for fruitful discussions about chemotaxis.

## Supporting Information

**Table S1.** Single *E. coli* persister cell waking by Ala. Single persister cells were observed using light microscopy (Zeiss Axio Scope.A1). The total number of *E. coli* BW25113 persister cells that wake on M9 5X Ala, M9 5X Asn, and no amino acid (aa) gel pads is shown after 6 h at 37 °C. Also, growth of exponential cells on gel pads that lack amino acids is shown after 6 h at 37 °C. 5X Ala and 5X Asn is 375 µg/mL. Total waking cells indicates the number of dividing or elongating cells. These results are the combined observations from two independent experiments. The microscope images are shown in **Fig. 1**.

|  | **Total cells** | **Dividing cells** | **Elongating cells** | **Total waking cells** | **% waking** | **Fold-change** |
| --- | --- | --- | --- | --- | --- | --- |
| **Ala** | 51 | 7 | 2 | 9 | 18 ± 1 | 9 |
| **Asn** | 46 | 1 | 0 | 1 | 2 ± 2 | 1 |
| **no aa** | 56 | 0 | 0 | 0 | 0 ± 0 | -∞ |
| **no aa (expon. cells)** | 104 | 12 | 0 | 12 | 12 ± 4 | - |

**Table S2.** Single persister cells wake primarily by sensing nutrients rather than spontaneously. Single persister cells were observed using light microscopy (Zeiss Axio Scope.A1). The total number of *E. coli* BW25113 wild type persister cells on the NaCl, NaCl + Amp, and NaCl + Amp + Ala agarose gel pads is shown after 6 h at 37 °C. Dead cells were identified by their disappearance. The concentration of NaCl was 0.85 %, Amp was 100 µg/mL, and Ala was 375 µg/mL. Fold-changes was based on NaCl + Amp. These results are the combined observations from two independent experiments. The microscope images are shown in **Fig. 2**.

|  | **Total cells** | **Dead cells** | **% of dead cells** | **Fold-change** |
| --- | --- | --- | --- | --- |
| **NaCl** | 144 | 0 | 0 ± 0 | - |
| **NaCl + Amp** | 164 | 13 | 7.5 ± 0.7 | 1 |
| **NaCl + Amp + Ala** | 135 | 47 | 33 ± 6 | 4.4 |

**Table S3.**
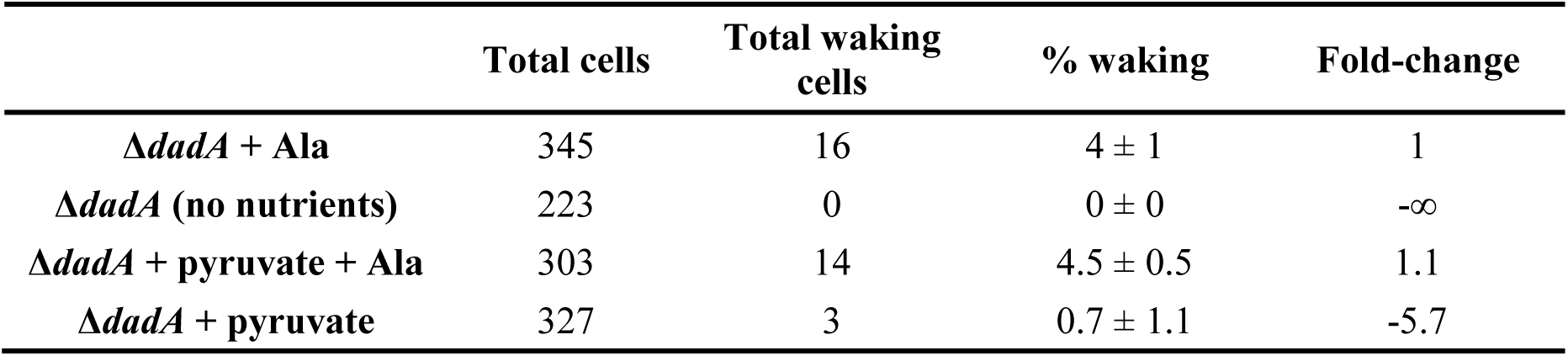
Single persister cells wake with alanine as a signal. Single persister cells were observed using light microscopy (Zeiss Axio Scope.A1). The total number and waking number of persister cells of BW25113 Δ*dadA* on M9 10X Ala, no nutrients, pyruvate (final conc. 0.24%) + 10 X Ala, and pyruvate (final conc. 0.24%) agarose gel pads are shown after 6 h at 37°C. These results are the combined observations from two independent experiments. The microscope images are shown in **Supplementary Figure 3**.

|  | Total cells | Total waking cells | % waking | Fold-change |
| --- | --- | --- | --- | --- |
| $\Delta dadA$ + Ala | 345 | 16 | $4 \pm 1$ | 1 |
| $\Delta dadA$ (no nutrients) | 223 | 0 | $0 \pm 0$ | $-\infty$ |
| $\Delta dadA$ + pyruvate + Ala | 303 | 14 | $4.5 \pm 0.5$ | 1.1 |
| $\Delta dadA$ + pyruvate | 327 | 3 | $0.7 \pm 1.1$ | -5.7 |

**Table S4.** Enhanced single persister cell waking on Ala via proteins identified by the ASKA library search. Single persister cells were observed using light microscopy (Zeiss Axio Scope.A1). The total number and waking number of persister cells of BW25113 producing the indicated proteins (PsiF, PanD, YmgF, YjcF, PptA, and CheY) are shown. Fold-change in waking is relative to BW25113 with the empty plasmid pCA24N. These results are the combined observations from two independent experiments after 18 hours. The microscope images are shown in **Supplementary Figure 5**.

|  | **Total**<br>**cells** | **Waking**<br>**cells** | **%**<br>**waking** | **Fold-**<br>**change** |
| --- | --- | --- | --- | --- |
| **Empty** | 103 | 2 | 3 ± 5 | 1 |
| **PsiF** | 91 | 10 | 11 ± 3 | 6.1 |
| **PanD** | 105 | 43 | 40 ± 30 | 22.8 |
| **YmgF** | 87 | 28 | 30 ± 10 | 17.9 |
| **YjcF** | 120 | 14 | 11.7 ± 0.3 | 6.5 |
| **PptA** | 118 | 15 | 8 ± 6 | 7.1 |
| **CheY** | 93 | 93 | 100 ± 0 | 33 |

**Table S5.**
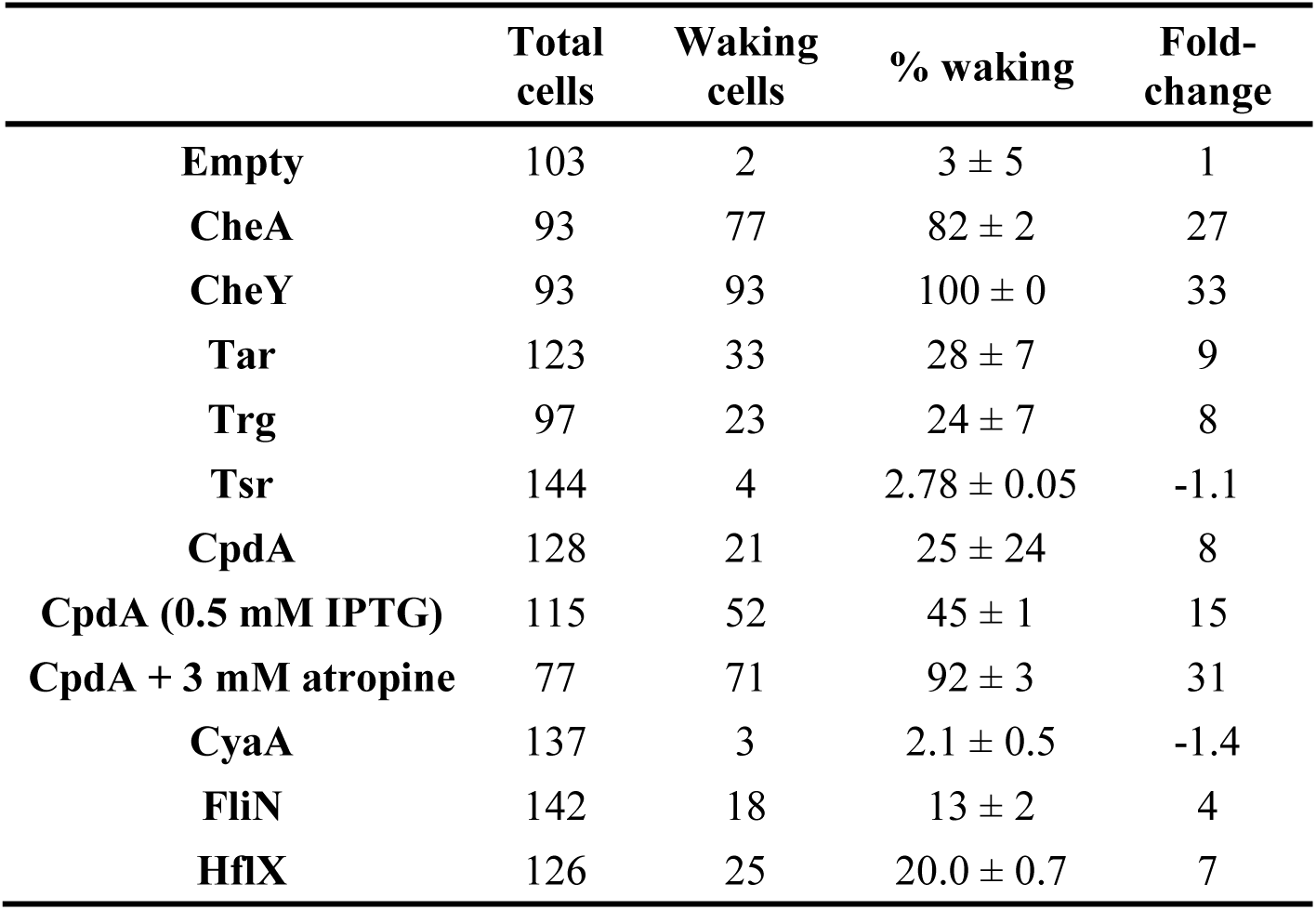
Single persister cell waking on Ala after producing chemotaxis-related, cAMP-related, and ribosome-resuscitation-related proteins. Single persister cells were observed using light microscopy (Zeiss Axio Scope.A1). The total number and waking number of persister cells are shown. Fold-change in waking is relative to BW25113 with the empty plasmid pCA24N (empty vector data from **Table S4**). IPTG was used only for producing CpdA as indicted, otherwise, protein production was from the leaky promoter. M9 5X Ala with 3 mM atropine in the agarose gel pad was used with cells producing CpdA. These results are the combined observations from two independent experiments after 18 hours. The microscope images are shown in **Fig. 3**.

**Table S6.** Persister cell formation on LB. Persister cells were observed on LB plates after 1 day, and the percentage of persister cell formation indicates the number of persisters formed relative to the BW25113 wild-type strain. “+” indicates roughly the same growth of the strain compared with the wild type. These results are the combined observations from two independent experiments.

|  | % persister formation | Growth |
| --- | --- | --- |
| **Wild type** | 100.0 ± 0.1 | + |
| *ΔarfB* | 98.9 ± 0.9 | + |
| *ΔcheA* | 99 ± 1 | + |
| *ΔcheY* | 101.6 ± 0.1 | + |
| *ΔcpdA* | 100.8 ± 0.3 | + |
| *Δcrp* | 102 ± 1 | + |
| *ΔcyaA* | 93.9 ± 2.4 | + |
| *ΔdadA* | 97 ± 4 | + |
| *ΔflgE* | 101.2 ± 0.6 | + |
| *ΔflhD* | 101.1 ± 0.2 | + |
| *ΔfliN* | 98.1 ± 0.8 | + |
| *ΔhflX* | 100.4 ± 0.2 | + |
| *ΔmotB* | 100.7 ± 0.4 | + |
| *ΔmotA* | 99.1 ± 0.2 | + |
| *ΔssrA'* | 101 ± 2 | + |
| *Δtar* | 98.3 ± 0.1 | + |
| *Δtrg* | 98.7 ± 0.4 | + |
| *ΔptsG* | 97 ± 2 | + |
| *Δcrr* | 97 ± 1 | + |
| *ΔmglB* | 98 ± 2 | + |
| *ΔmalE* | 100.8 ± 0.4 | + |
| *ΔsmpB* | 96 ± 2 | + |
| **pCA24N** | 100.0 ± 0.5 | + |
| **ArfB** | 97.4 ± 0.7 | + |
| **CheA** | 95.6 ± 0.6 | + |
| **CheY** | 99.6 ± 0.5 | + |
| **CpdA** | 99.7 ± 0.5 | + |
| **Crp** | 98.2 ± 0.7 | + |
| **CyaA** | 99.7 ± 0.7 | + |
| **FlhD** | 98.0 ± 0.7 | + |
| **FliN** | 97.6 ± 0.6 | + |
| **HflX** | 97.7 ± 0.9 | + |
| **Tar** | 98.8 ± 0.8 | + |
| **Trg** | 97.9 ± 0.7 | + |
| **PtsG** | 97 ± 1 | + |
| **Crr** | 93.6 ± 0.2 | + |

**Table S7.** Single persister cell waking on Ala after inactivating flagellar proteins MotB, FhlD, and FliN. Single persister cells were observed using light microscopy (Zeiss Axio Scope.A1). The total number and waking number of persister cells are shown. Fold-change in waking is relative to wild-type BW25113 (wild-type data from **Table S1**). These results are the combined observations from two independent experiments after 6 hours. The microscope images are shown in **Supplementary Figure 10**.

|  | **Total**<br>**cells** | **Waking**<br>**cells** | **%**<br>**waking** | **Fold-**<br>**change** |
| --- | --- | --- | --- | --- |
| **Wild type** | 51 | 9 | 18 ± 1 | 1 |
| *ΔmotB* | 57 | 17 | 30 ± 10 | 1.7 |
| *ΔflhD* | 81 | 5 | 6.1 ± 0.4 | -3 |
| *ΔfliN* | 96 | 5 | 5 ± 2 | -3.6 |

**Table S8.**
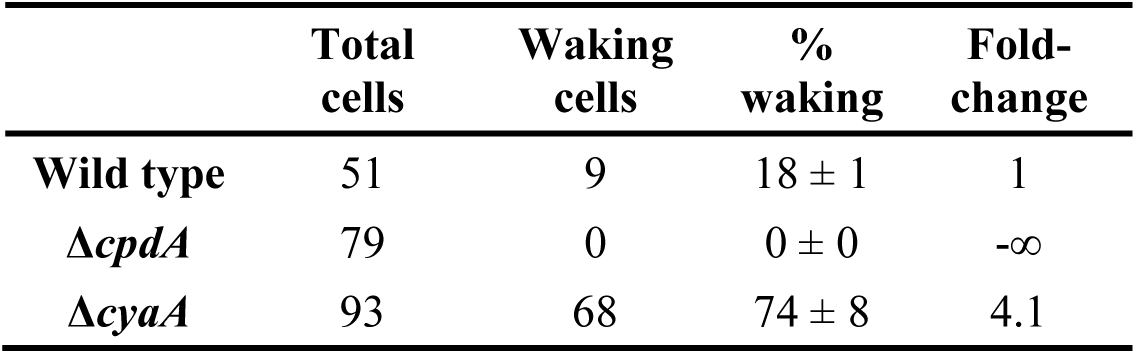
Single persister cell waking on Ala after inactivating CpdA and CyaA. Single persister cells were observed using light microscopy (Zeiss Axio Scope.A1). The total number and waking number of persister cells are shown. Fold-change in waking is relative to wild-type BW25113 (wild-type data from **Table S1**). These results are the combined observations from two independent experiments after 6 hours. The microscope images are shown in **Supplementary Figure 13**.

|  | **Total**<br>**cells** | **Waking**<br>**cells** | **%**<br>**waking** | **Fold-**<br>**change** |
| --- | --- | --- | --- | --- |
| **Wild type** | 51 | 9 | $18 \pm 1$ | 1 |
| ***ΔcpdA*** | 79 | 0 | $0 \pm 0$ | $-\infty$ |
| ***ΔcyaA*** | 93 | 68 | $74 \pm 8$ | 4.1 |

**Table S9.** Single persister cell waking on Ala after inactivating ribosome rescue proteins HflX and ArfB and inactivating trans-translation (SsrA). Single persister cells were observed using light microscopy (Zeiss Axio Scope.A1). The total number and waking number of persister cells are shown. Fold-change in waking is relative to wild-type BW25113 (wild-type data from Table S1). These results are the combined observations from two independent experiments after 6 hours. The microscope images are shown in **Supplementary Figure 16**.

|  | **Total**<br>**cells** | **Waking**<br>**cells** | **%**<br>**waking** | **Fold-**<br>**change** |
| --- | --- | --- | --- | --- |
| **Wild type** | 51 | 9 | 18 ± 1 | 1 |
| ***ΔhflX*** | 96 | 0 | 0 ± 0 | -∞ |
| ***ΔarfB*** | 101 | 15 | 15 ± 2 | -1.2 |
| ***ΔssrA*** | 123 | 1 | 0.8 ± 1.1 | -22.5 |

**Table S10.** Single persister cell waking on glucose or alanine gel pads. Single wild-type persister cells were observed using light microscopy (Zeiss Axio Scope.A1). The total number and waking number of persister cells on minimal medium with 0.4% glucose and 0.04% alanine are shown (0.4% glucose inhibited resuscitation with Ala). Fold-change in waking is relative to alanine (wild-type Ala data from **Table S1**). These results are the combined observations from two independent experiments after 6 hours. The microscope images are shown in **Figure 4**.

|  | **Glucose** | **Alanine** |
| --- | --- | --- |
| **Total cells** | 354 | 51 |
| **Waking cells** | 326 | 9 |
| **Waking %** | 92 ± 2 | 18 ± 1 |
| **Fold-change** | 5.1 | 1 |

**Table S11.** Single persister cell waking on glucose gel pads. Single persister cells were observed using light microscopy (Zeiss Axio Scope.A1). The total number and waking number of persister cells are shown. Fold-change in waking is relative to wild-type BW25113 (same field of view with 354 cells as in **Table S10** but cells wake here for only 3 h vs. 6 h in **Table S10**). These results are the combined observations from two independent experiments after 3 hours. The microscope images are shown in **Figure 5**.

|  | Total cells | Waking cells | % waking | Fold-change |
| --- | --- | --- | --- | --- |
| wild type | 354 | 74 | 22 ± 3 | 1 |
| *Δtrg* | 283 | 12 | 5 ± 1 | -4.6 |
| *ΔptsG* | 212 | 1 | 0.3 ± 0.5 | -69.2 |
| *Δcrr* | 369 | 44 | 11±3 | -2 |
| *ΔmglB* | 311 | 55 | 17.8 ± 0.5 | -1.2 |
| *ΔmalE* | 329 | 77 | 23.3 ± 0.5 | 1.1 |
| *ΔcpdA* | 305 | 20 | 6 ± 2 | -3.7 |
| *ΔcyaA* | 279 | 139 | 50 ± 2 | 2.3 |
| *Δhfq* | 201 | 23 | 12 ± 1 | -1.9 |
| *ΔsmpB* | 352 | 132 | 35 ± 5 | 1.6 |
| *ΔssrA* | 316 | 22 | 6.9 ± 0.1 | -3.2 |

**Table S12.** Single persister cell waking on glucose gel pads. Single persister cells were observed using light microscopy (Zeiss Axio Scope.A1). The total number and waking number of persister cells are shown. Fold-change in waking is relative to BW25113 with the empty plasmid pCA24N. These results are the combined observations from two independent experiments after 5 hours. The microscope images are shown in **Figure 6**.

|  | **Total cells** | **Waking cells** | **% waking** | **Fold-change** |
| --- | --- | --- | --- | --- |
| **Empty** | 284 | 7 | $2.5 \pm 0.3$ | 1 |
| **PtsG** | 195 | 0 | $0 \pm 0$ | $-\infty$ |
| **Crr** | 356 | 5 | $1.4 \pm 0.2$ | -1.8 |
| **CheY** | 339 | 129 | $39 \pm 4$ | 15.9 |
| **CpdA** | 269 | 104 | $39 \pm 1$ | 15.9 |
| **HflX** | 321 | 11 | $3.41 \pm 0.07$ | 1.4 |
| **Hfq** | 207 | 11 | $5.0 \pm 0.9$ | 2 |

**Table S13.**
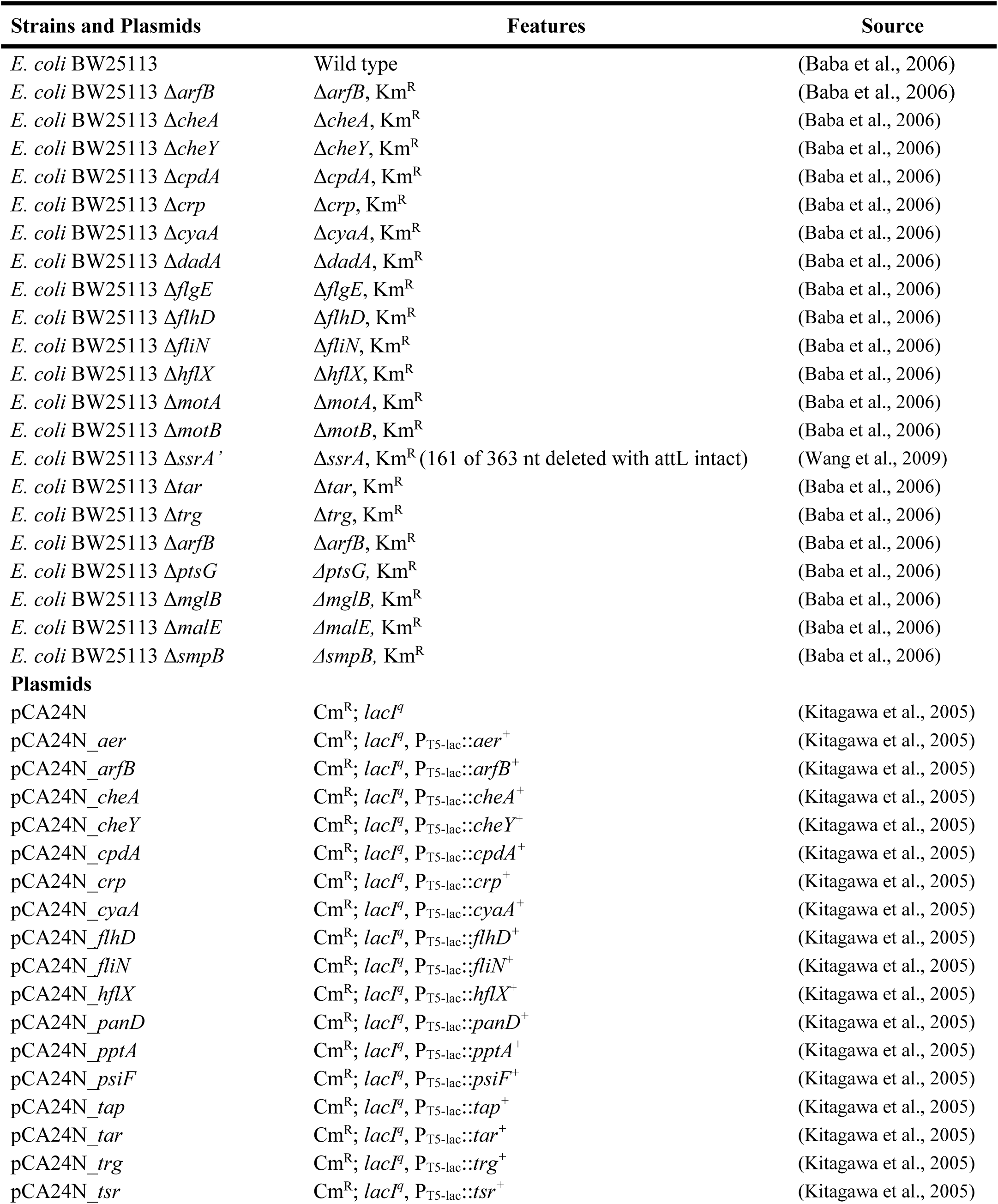
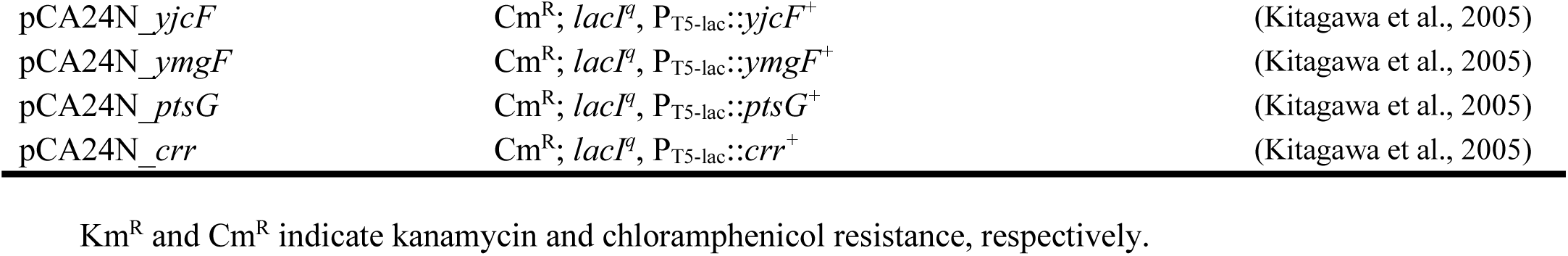
Bacterial strains and plasmids used in this study.

**Table S14.** Supplementation levels (1X) for amino acids in M9 minimal meium.

| No. | Amino acid | Final concentration<br>($\mu\text{g/mL}$) |
| --- | --- | --- |
| 1 | 1 % L-alanine | 75 |
| 2 | 2 % L-arginine | 145 |
| 3 | 1 % L-asparagine | 75 |
| 4 | 1 % L aspartic acid | 75 |
| 5 | 2 % L-cysteine | 50 |
| 6 | 2 % L-histidine | 42 |
| 7 | 1 % L-glutamic acid | 75 |
| 8 | 1 % L-glutamine | 75 |
| 9 | 2 % glycine | 110 |
| 10 | 1 % L-isoleucine | 42 |
| 11 | 1 % L-leucine | 41 |
| 12 | 1 % L-lysine | 75 |
| 13 | 2 % L-methionine | 25 |
| 14 | 1 % L-phenylalanine | 75 |
| 15 | 4 % L-proline | 164 |
| 16 | 2 % L-serine | 42 |
| 17 | 2 % L-threonine | 82 |
| 18 | 0.25 % L-tryptophan | 18 |
| 19 | 1 % L-tyrosine | 75 |
| 20 | 1 % L-valine | 42 |

**Supplementary Figure 1.**
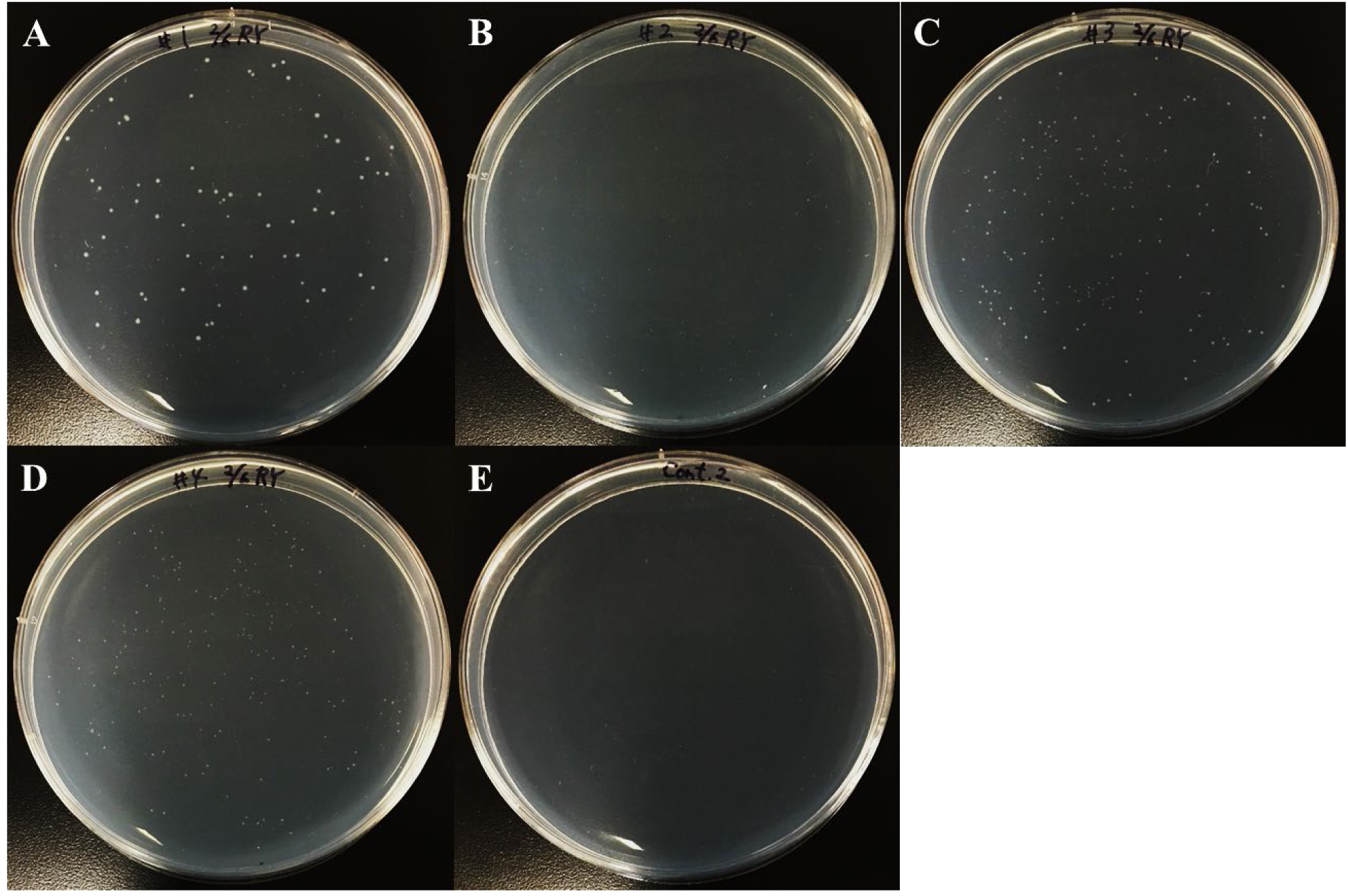
Persister cells waking on agar plates with groups of amino acids. *E. coli* BW25113 persister cells were incubated at 37 °C for 4 days on M9 agar plates with groups of five different amino acids: **(A) #1**: Arg, Ser, Cys, Ala, and Phe, **(B) #2**: His, Thr, Gly, Val, and Tyr, **(C) #3**: Lys, Asn, Pro, Ile, and Trp, **(D) #4**: Asp, Glu, Gln, Leu, and Met, and **(E)** no amino acids. These amino acids were used at 1X concentration (**Table S14**). One representative plate of two independent cultures is shown.

**Supplementary Figure 2.**
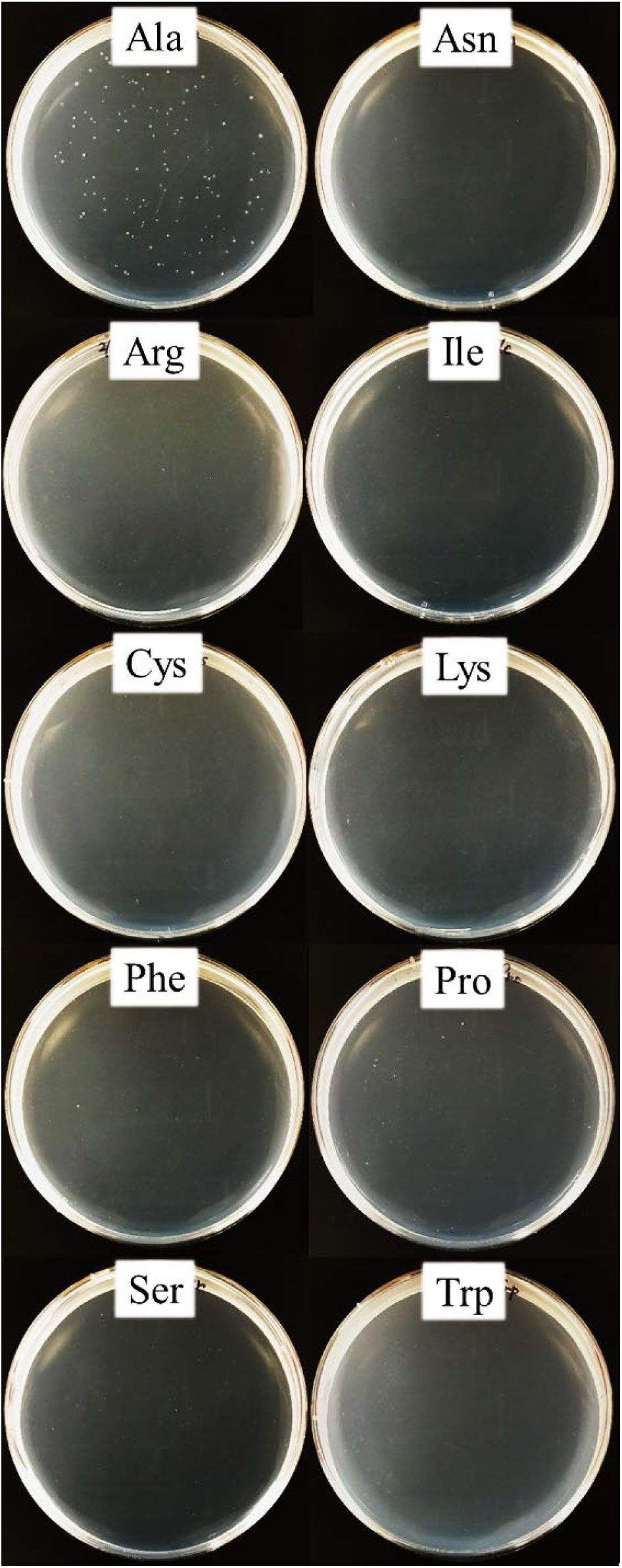
Persister cells waking on agar plates with alanine. *E. coli* BW25113 persister cells were incubated at 37 °C for 4 days on M9 agar plates with 10 different amino acids (Arg, Ser, Cys, Ala, Phe, Lys, Asn, Pro, Ile, and Trp). These amino acids were used at 1X concentration (**Table S14**). One representative plate of two independent cultures is shown.

**Supplementary Figure 3.**
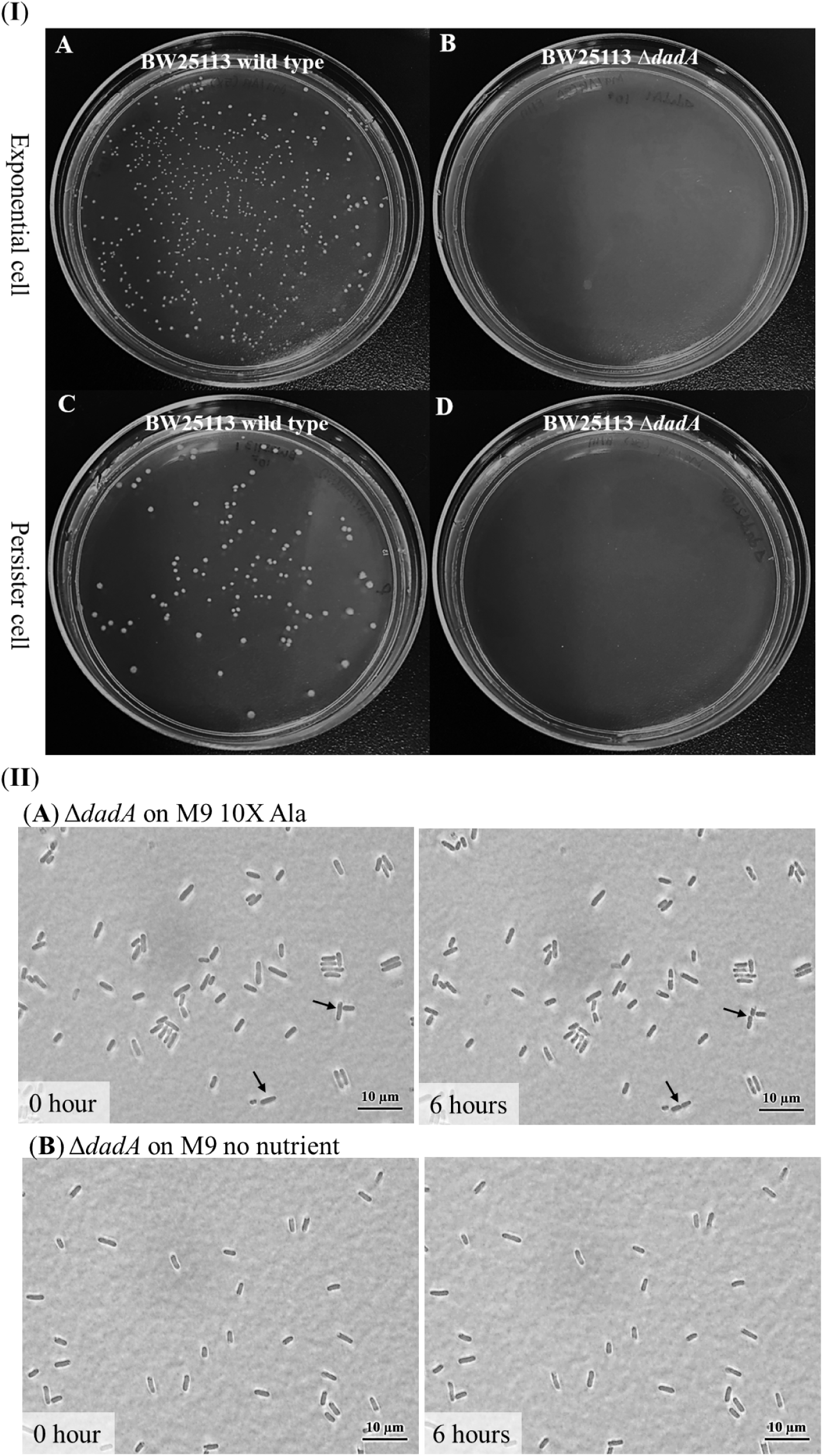
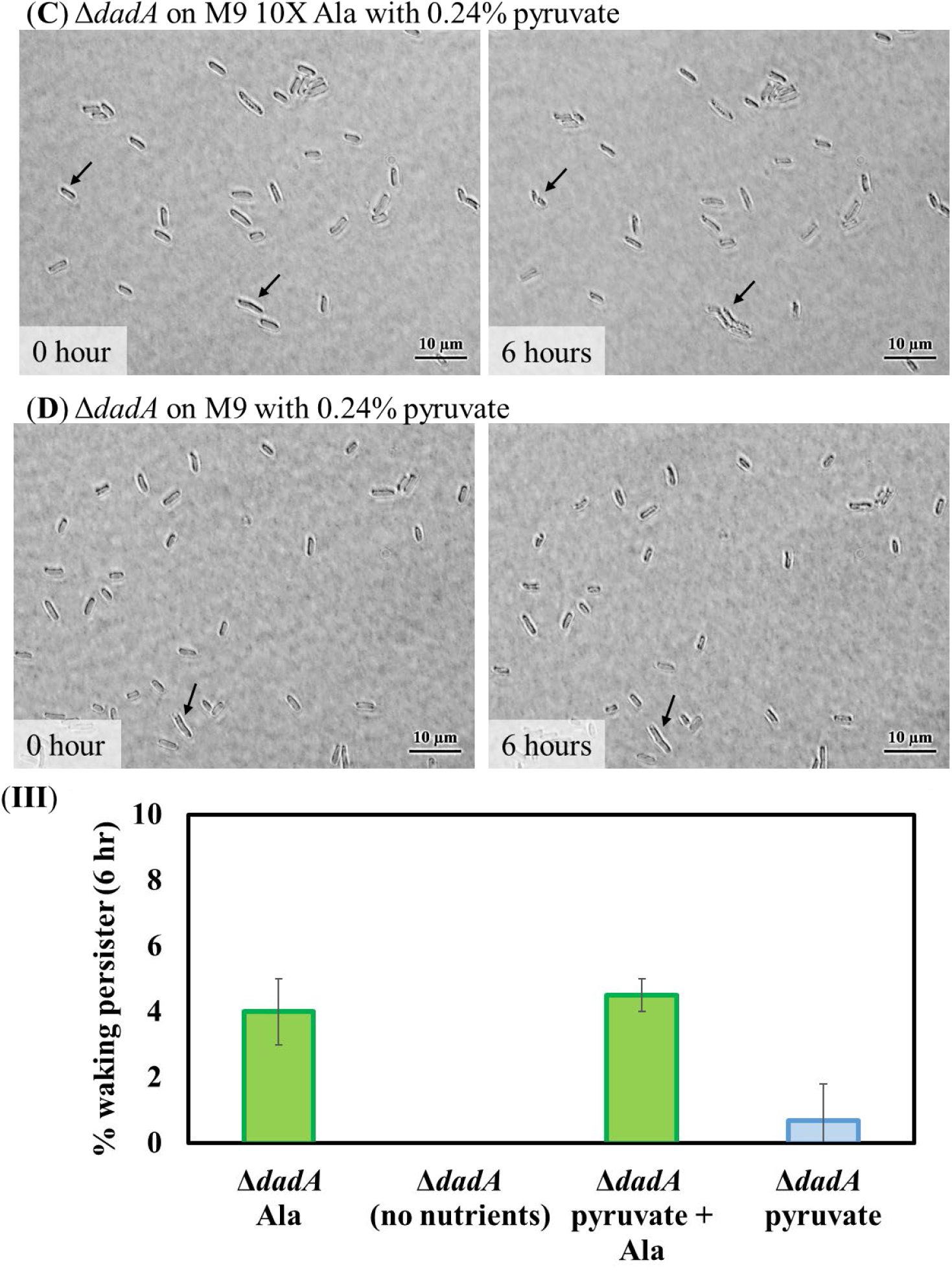
Alanine is a waking signal. (**I**) Exponential wild-type *E. coli* BW25113 (**A**), exponential BW25113 Δ*dadA* (**B**), persister wild-type *E. coli* BW25113 (**C**), and persister BW25113 Δ*dadA* (**D**) incubated at 37 °C on M9 5X Ala agar plates for 3 days. (**II**) Persister cells of BW25113 Δ*dadA* on M9 10X Ala gel pads (**A**), on M9 no nutrient gel pads (**B**), on M9 10X Ala with 0.24% pyruvate (**C**), and on M9 with 0.24% pyruvate (**D**) after 6 h at 37°C. Black arrows indicate cells that resuscitate. Scale bar indicates 10 µm. One representative plate of two independent cultures is shown. (**III**) *dadA* persister cell waking (%) after 6 hours. Tabulated cell numbers are shown in **Supplementary Table 3**.

**Supplementary Figure 4.**
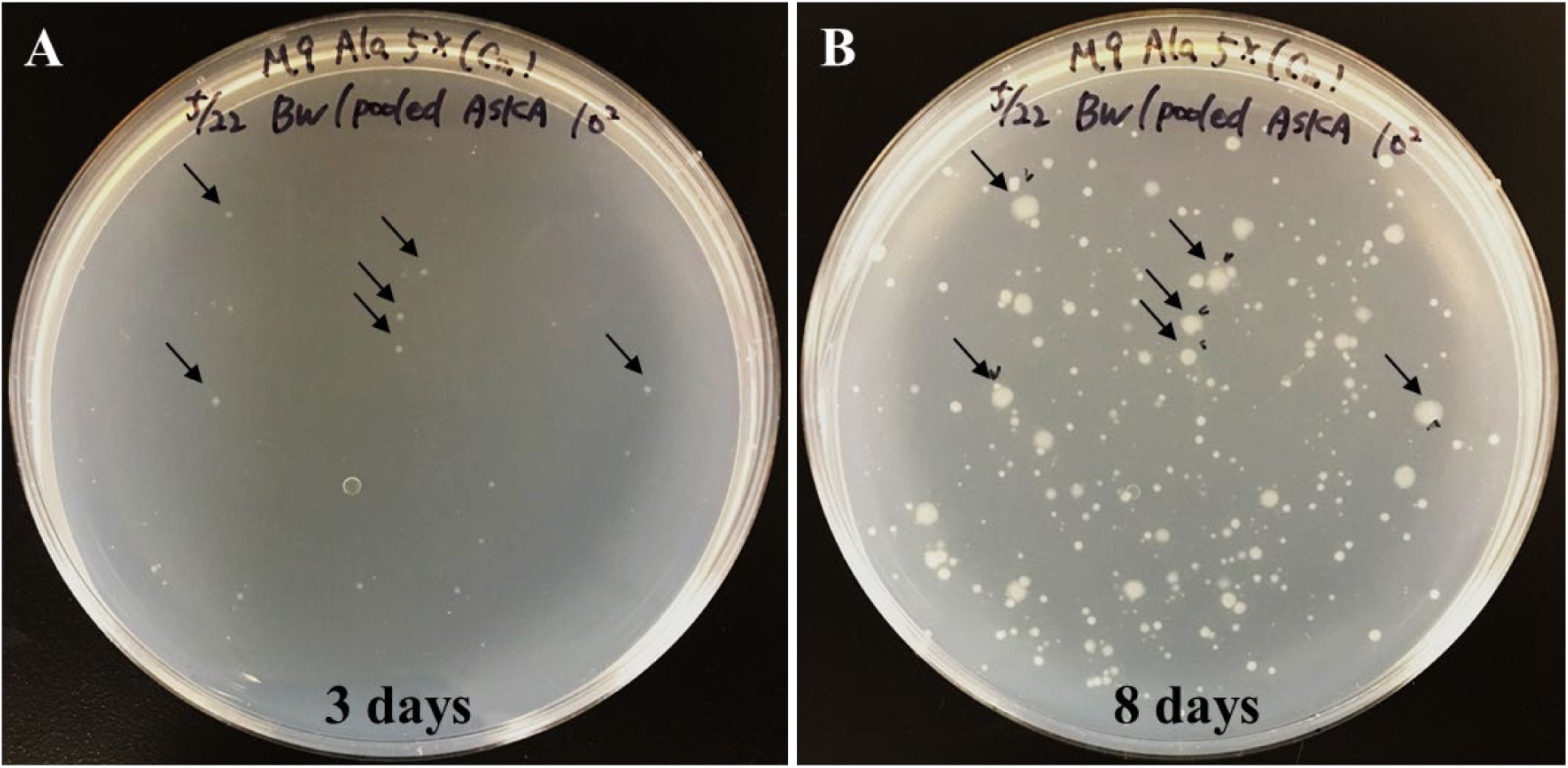
Screening persister cell waking with the pooled ASKA plasmids on Ala. *E. coli* BW25113 persisters containing each of the pooled ASKA plasmids were incubated at 37 °C on M9 5X Ala agar plates with 30 μg/mL Cm for (**A**) 3 days and (**B**) 8 days. Faster growing colonies are indicated with black arrows. Prior to forming persister cells, cultures were grown without IPTG with 30 µg/mL Cm. One representative plate of two independent cultures is shown.

**Supplementary Figure 5.**
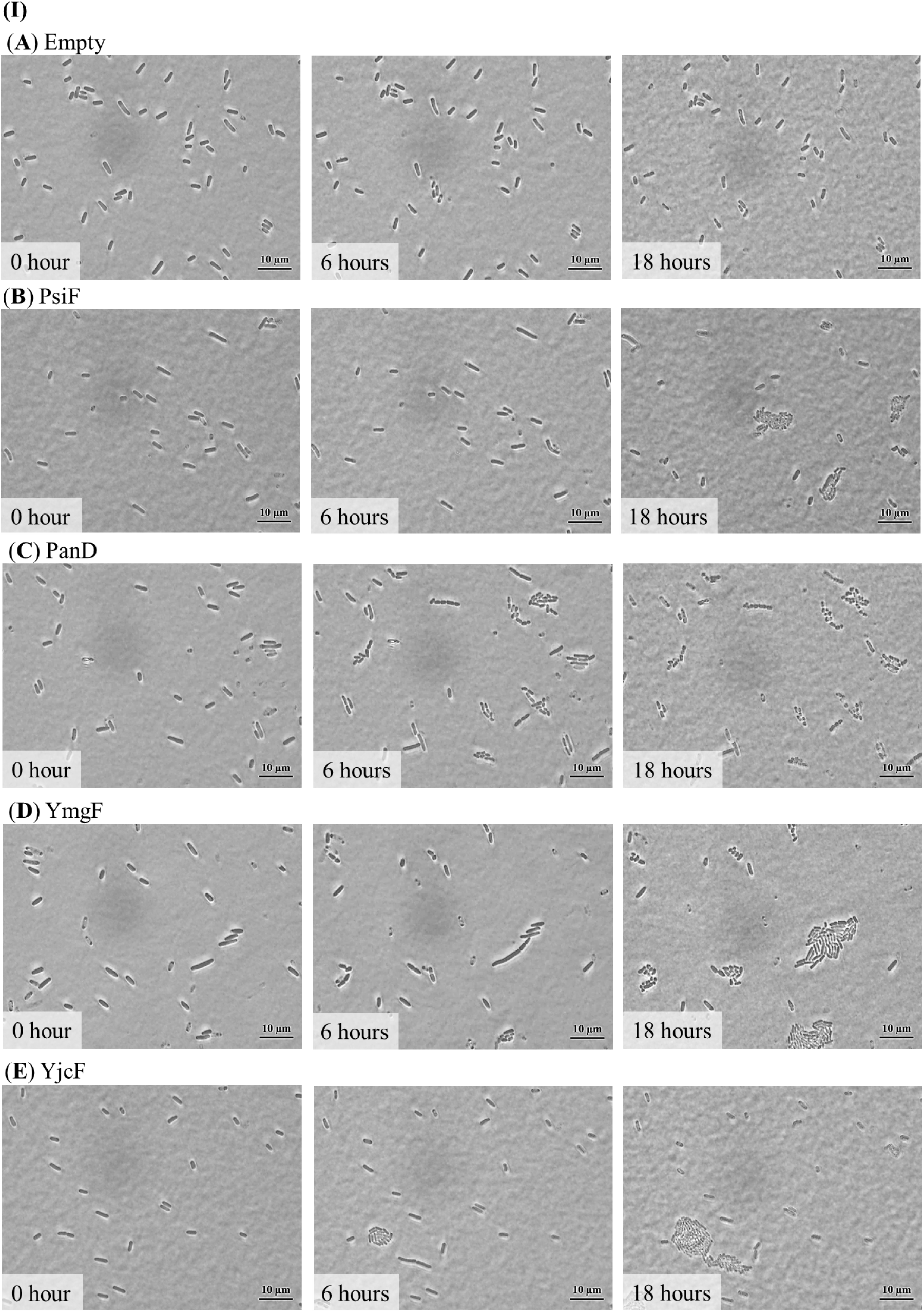
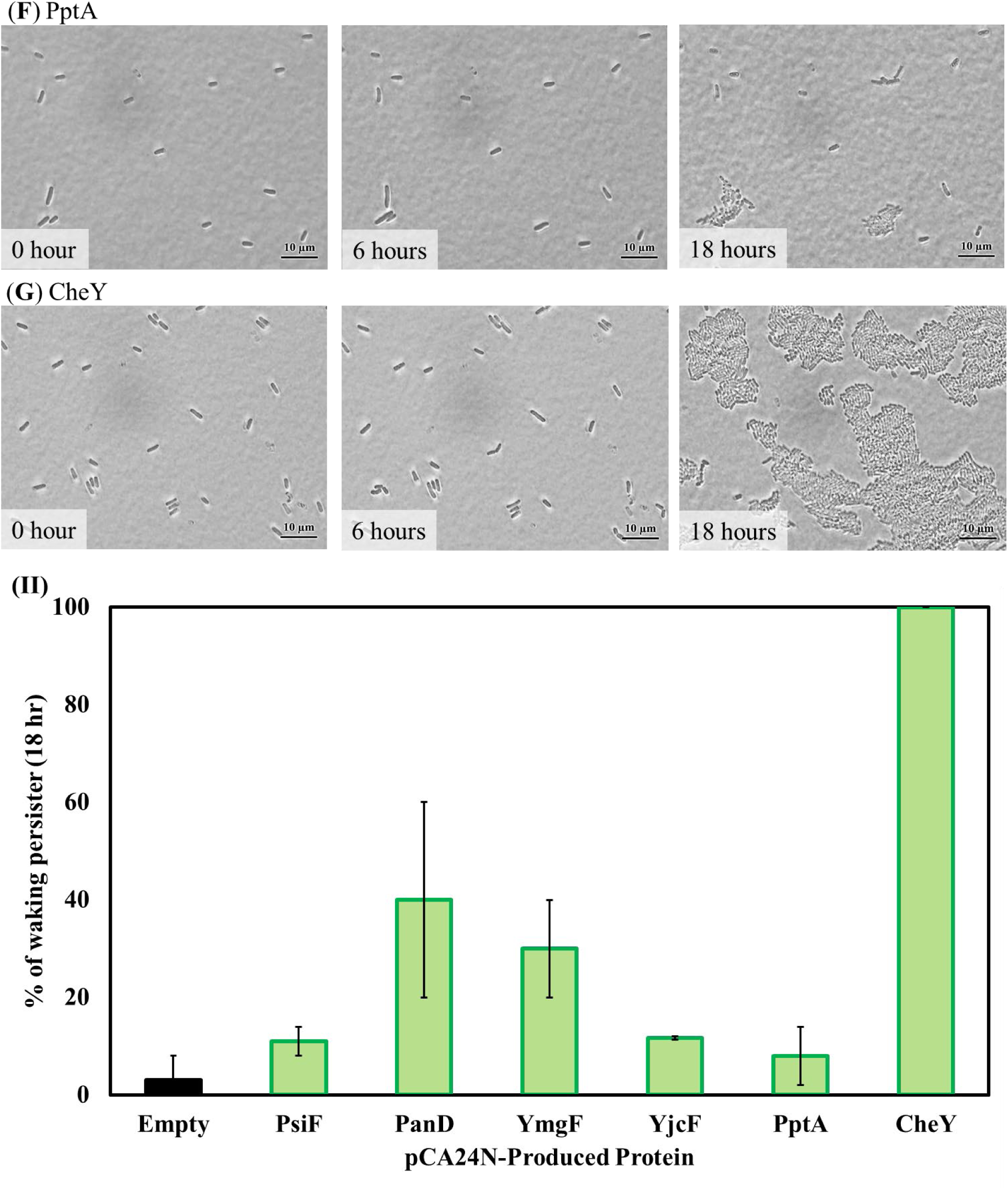
Single persister cell waking on Ala related to the ASKA screen. (**I**) *E. coli* BW25113 persister cells containing (**A**) pCA24N (empty plasmid control), (**B**), pCA24N_*psiF*, (**C**) pCA24N_*panD*, (**D**) pCA24N_*ymgF*, (**E**) pCA24N_*yjfF*, (**F**) pCA24N_*pptA*, and (**G**) pCA24N_*cheY* were incubated at 37°C on M9 5X Ala agarose gel pads for 18 h. Scale bar indicates 10 µm. Representative results from two independent cultures are shown. Prior to forming persister cells, cultures were grown without IPTG with 30 µg/mL Cm. (**II**) Persister cell waking (%) after 18 hours.

**Supplementary Figure 6.**
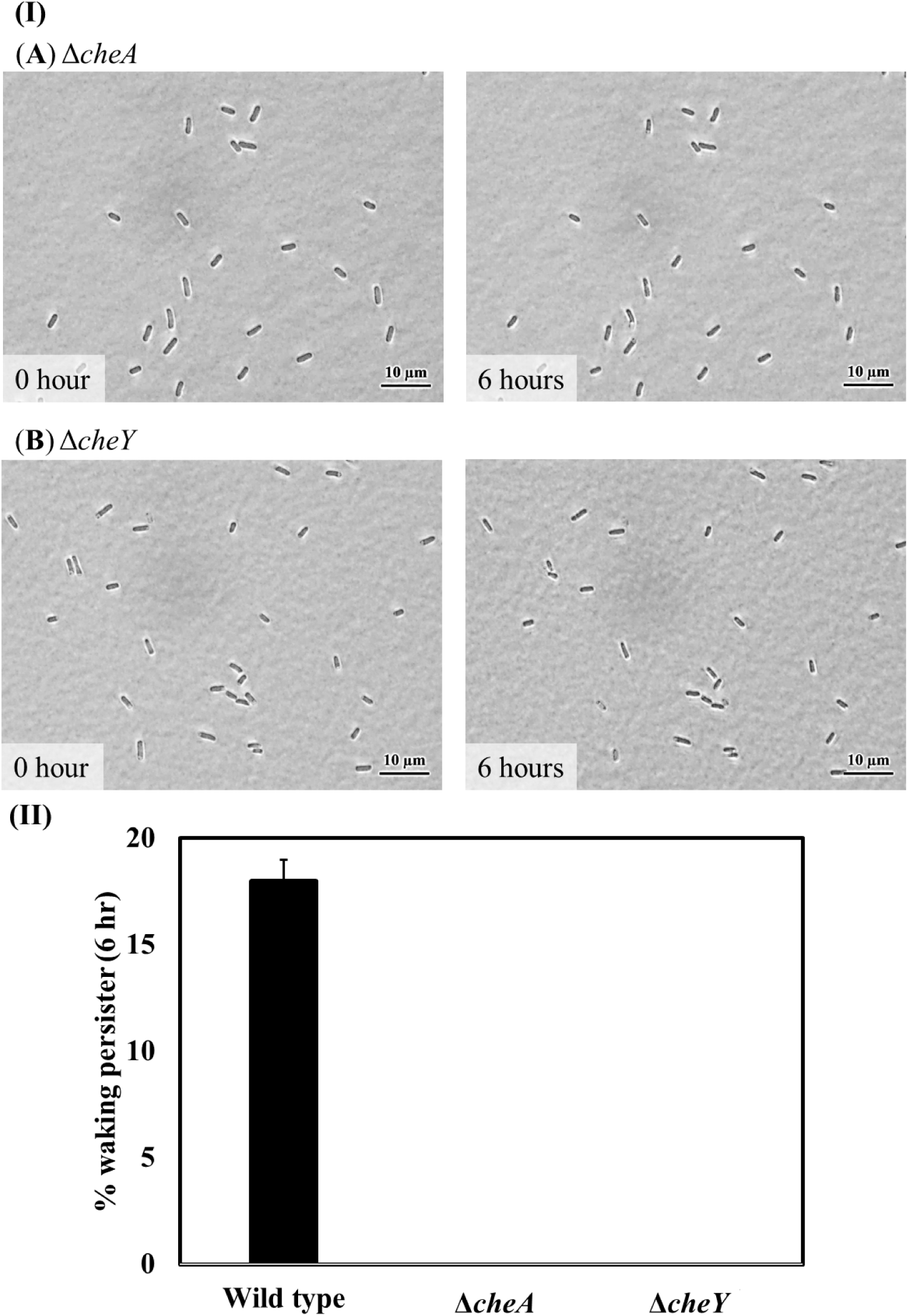
Single persister cell waking on Ala after inactivating chemotaxis proteins CheA and CheY. (**I**) Persister cells of (**A**) BW25113 Δ*cheA* and (**B**) BW25113 Δ*cheY* waking on M9 5X Ala agarose gel pads after 6 h at 37°C. Scale bar indicates 10 µm. Representative results from two independent cultures are shown. (**II**) Persister cell waking (%) after 6 hours.

**Supplementary Figure 7.**
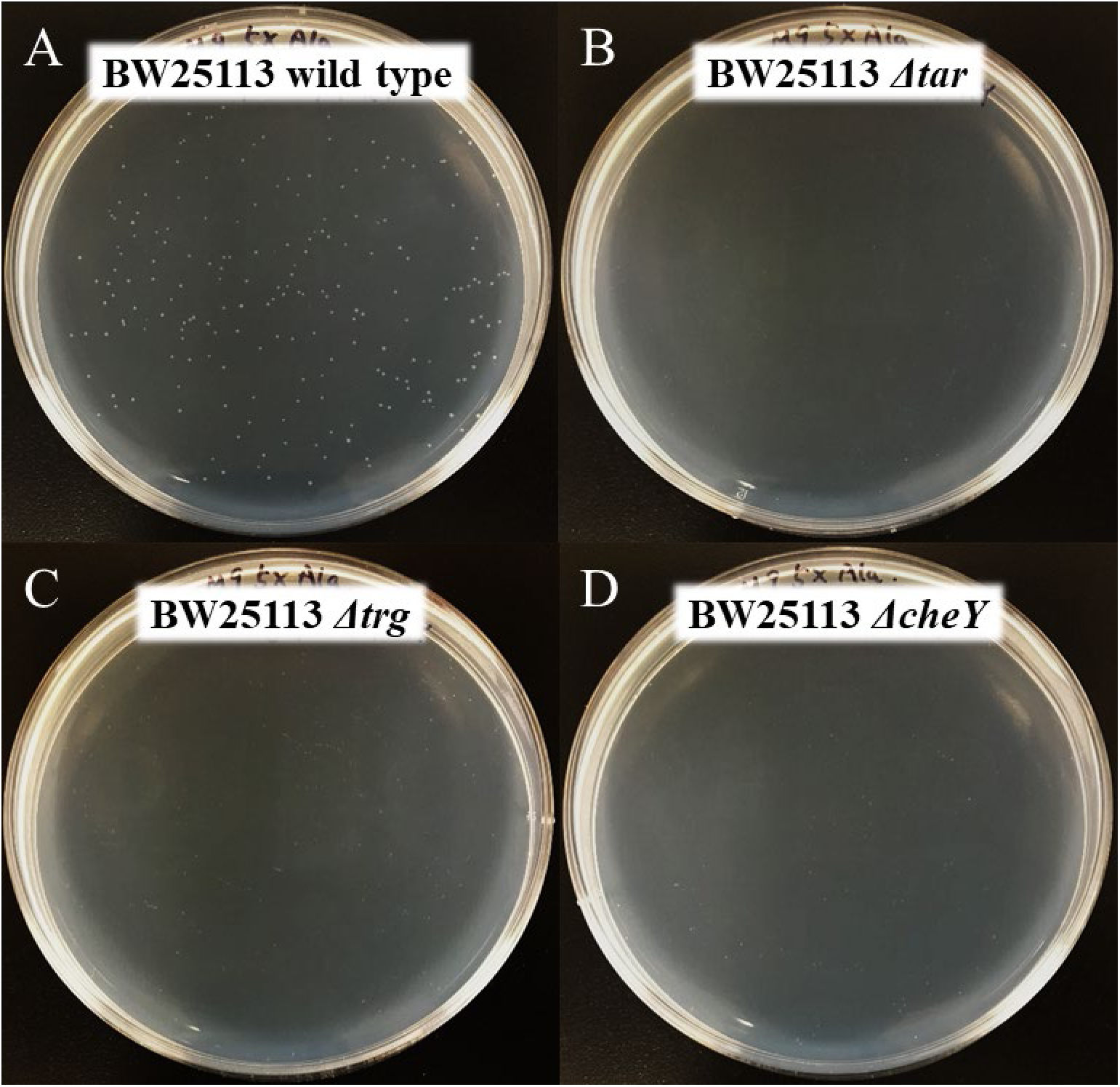
Persister cell waking on Ala agar plates after inactivating the chemotaxis-related proteins Tar, Trg, and CheY. Persister cells were incubated at 37 °C on M9 5X Ala agar plates for 2 days. (**A**) wild type BW25113, (**B**) BW25113 Δ*tar*, (**C**) BW25113 Δ*trg*, and (**D**) BW25113 Δ*cheY*. One representative plate of two independent cultures is shown.

**Supplementary Figure 8.**
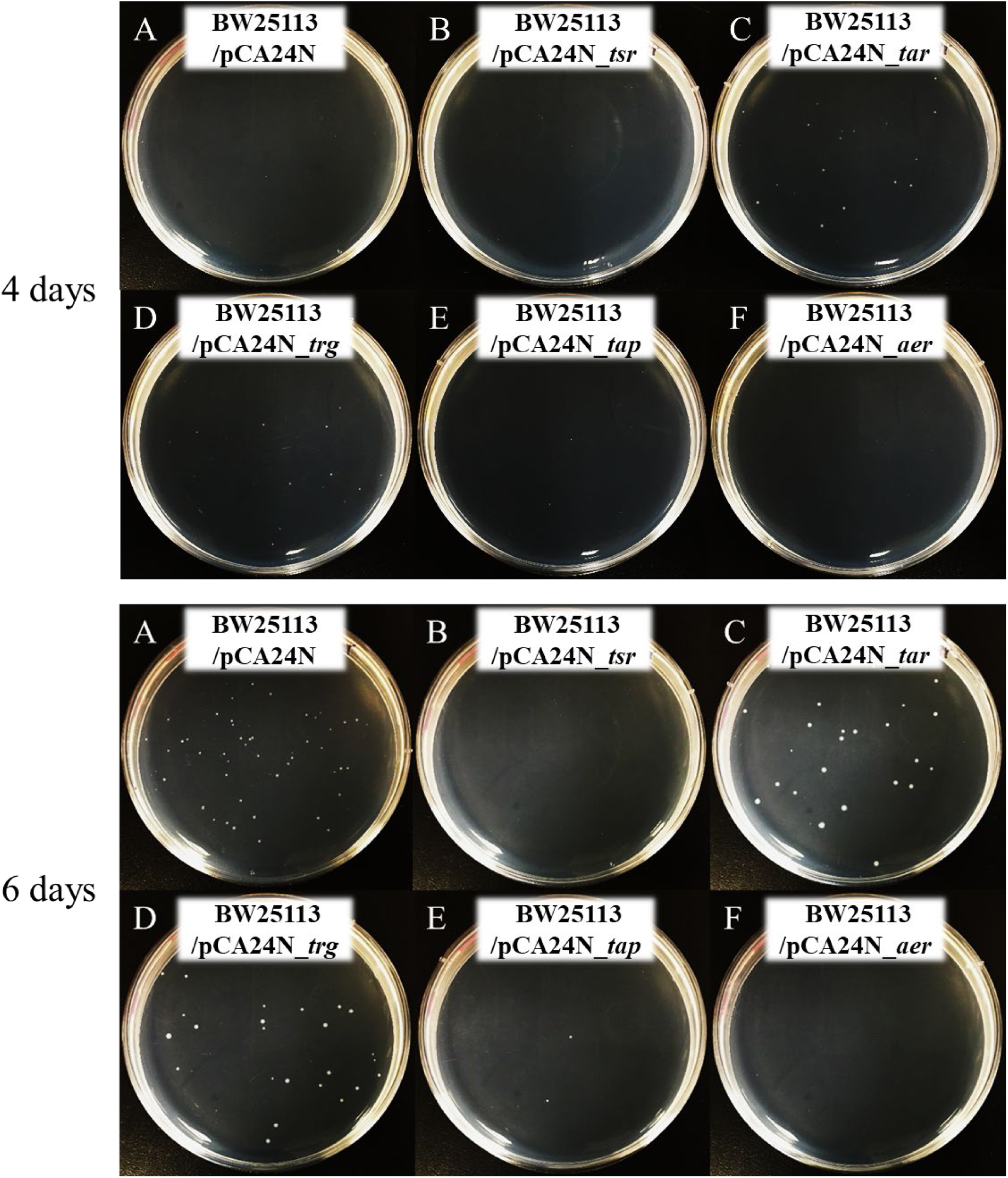
Persister cell waking on Ala agar plates after producing chemotaxis proteins. Persister waking after 4 days (upper panel) and after 6 days (lower panel) incubation at 37 °C on M9 5X Ala agar plates. (**A**) BW25113/pCA24N, (**B**) BW25113/pCA24N_*tsr*, (**C**) BW25113/pCA24N_*tar*, (**D**) BW25113/pCA24N_*trg*, (**E**) BW25113/pCA24N_*tap*, and (**F**) BW25113/pCA24N_*aer.* Prior to forming persister cells, cultures were grown without IPTG with 30 µg/mL Cm. One representative plate of two independent cultures is shown.

**Supplementary Figure 9.**
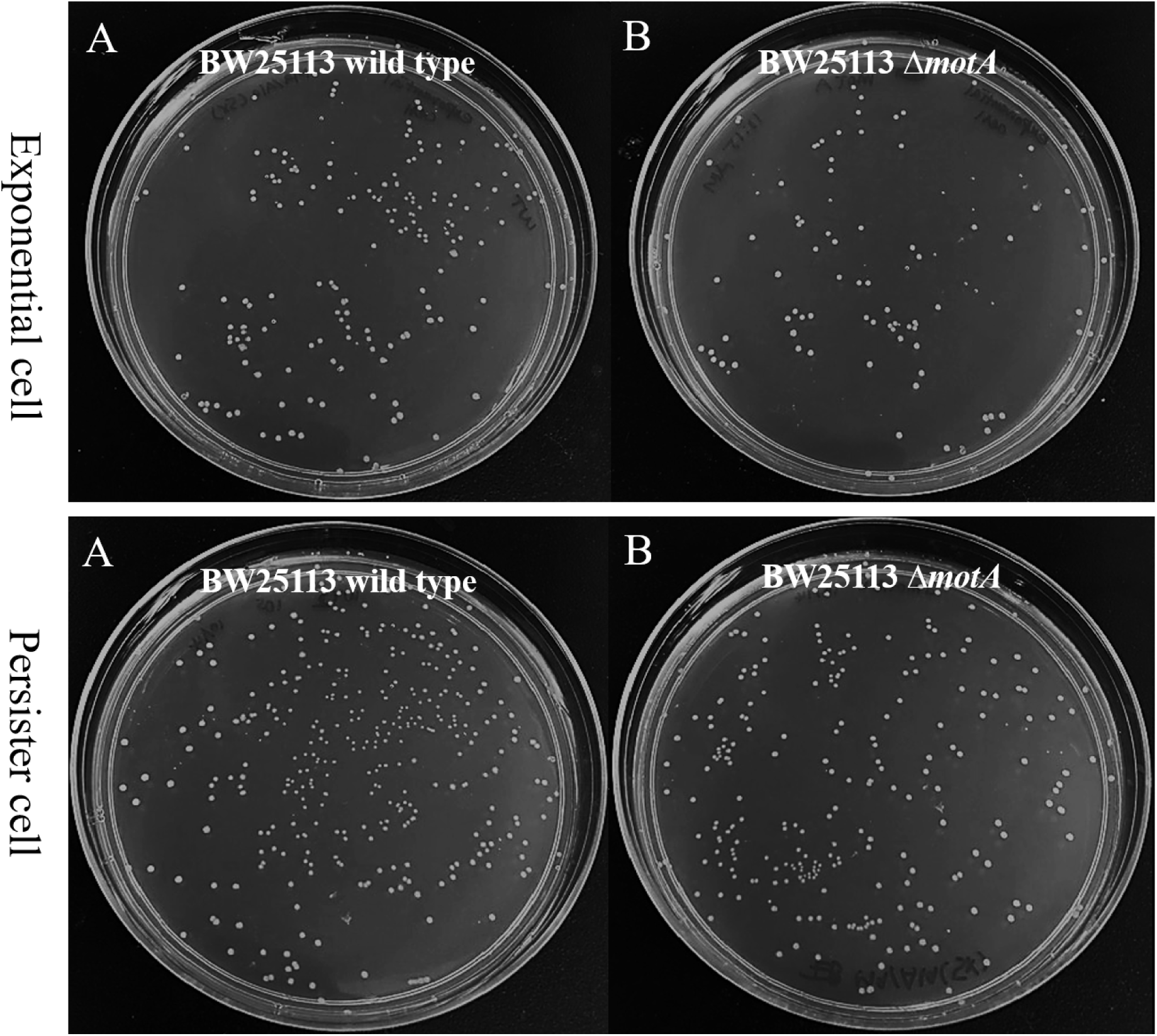
Persister cell waking on Ala agar plates after inactivating flagellar motor complex proteins MotA. Cells were incubated at 37 °C on M9 5X Ala agar plates for 3 days. Upper panel: Exponential cells (**A**) wild type BW25113 and (**B**) BW25113 Δ*motA*. Lower panel: Persister cells of (**A**) wild type BW25113 and (**B**) BW25113 Δ*motA*. One representative plate of two independent cultures is shown.

**Supplementary Figure 10.**
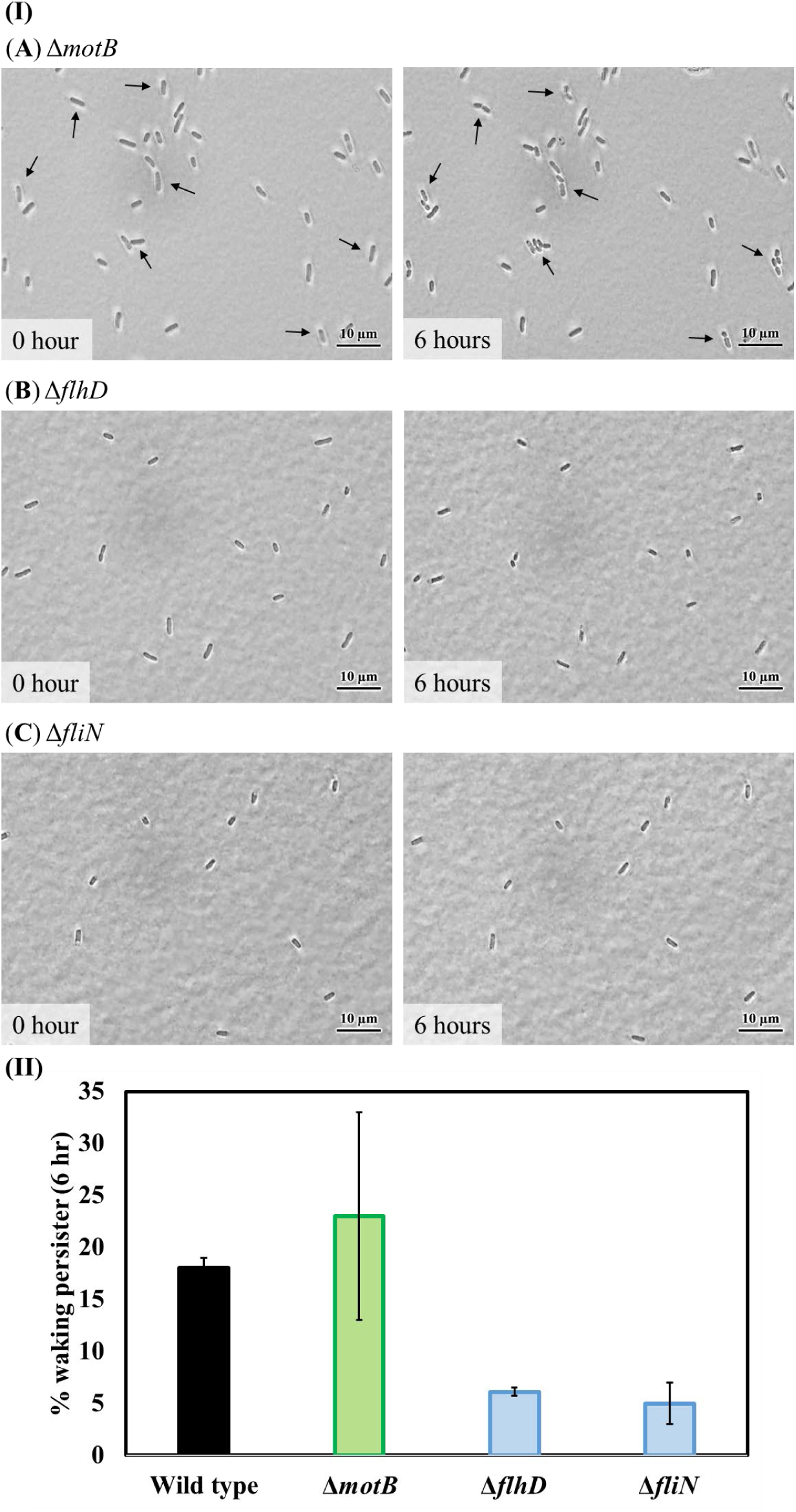
Single persister cell waking on Ala after inactivating flagellar proteins MotB, FhlD, and FliN. (**I**) Persister cells of (**A**) BW25113 Δ*motB*, (**B**) BW25113 Δ*flhD*, and (**C**) BW25113 Δ*fliN* waking on M9 5X Ala agarose gel pads after 6 h at 37°C. Black arrows indicate cells that resuscitate. Scale bar indicates 10 µm. Representative results from two independent cultures are shown. (**II**) Persister cell waking (%) after 6 hours.

**Supplementary Figure 11.**
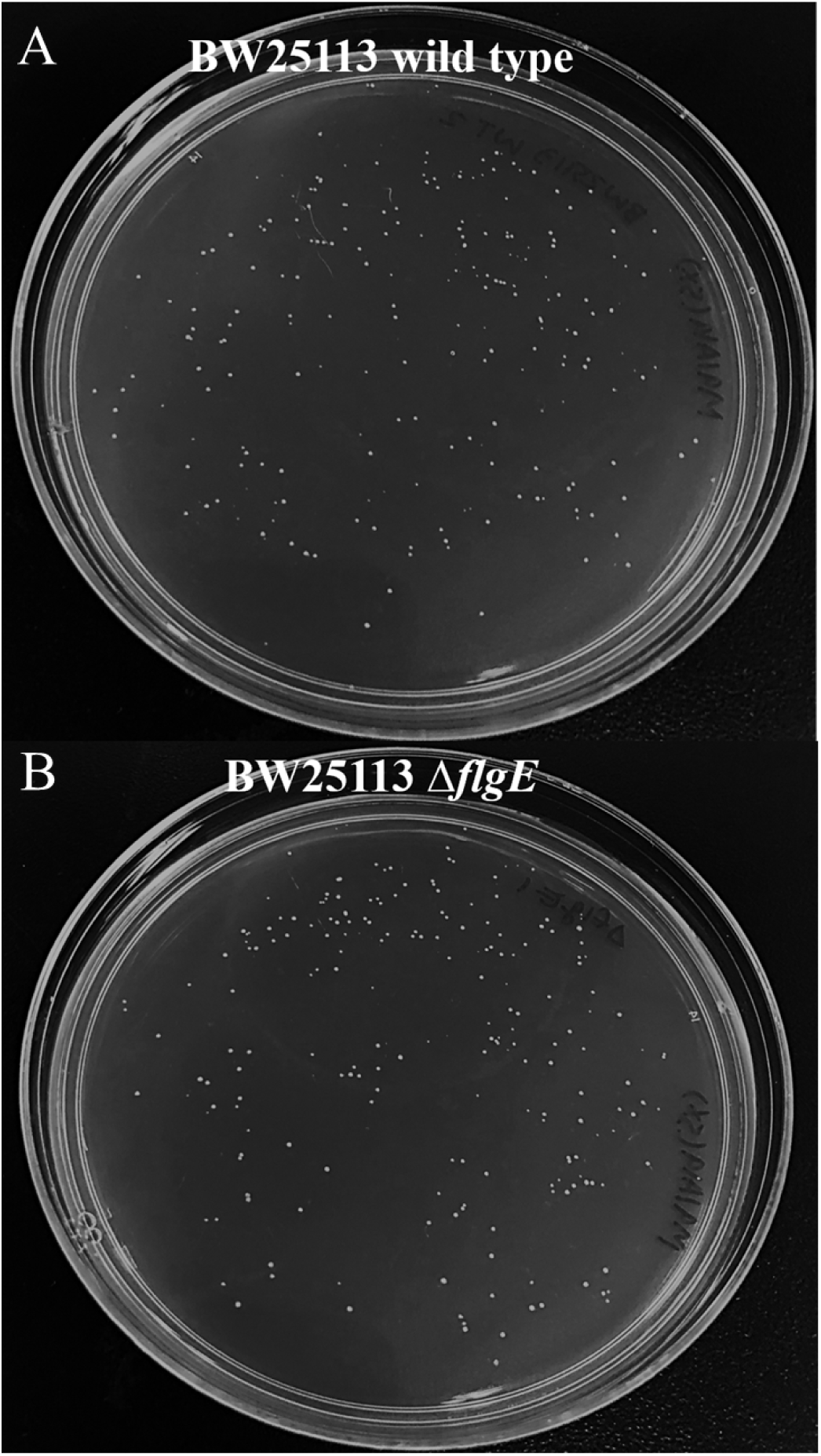
Persister cell waking on Ala agar plates after inactivating the FlgE flagellum hook protein. *E. coli* BW25113 persister cells of (**A**) wild type BW25113 and (**B**) BW25113 Δ*flgE* were incubated at 37 °C on M9 5X Ala agar plates for 2 days. One representative plate of two independent cultures is shown.

**Supplementary Figure 12.**
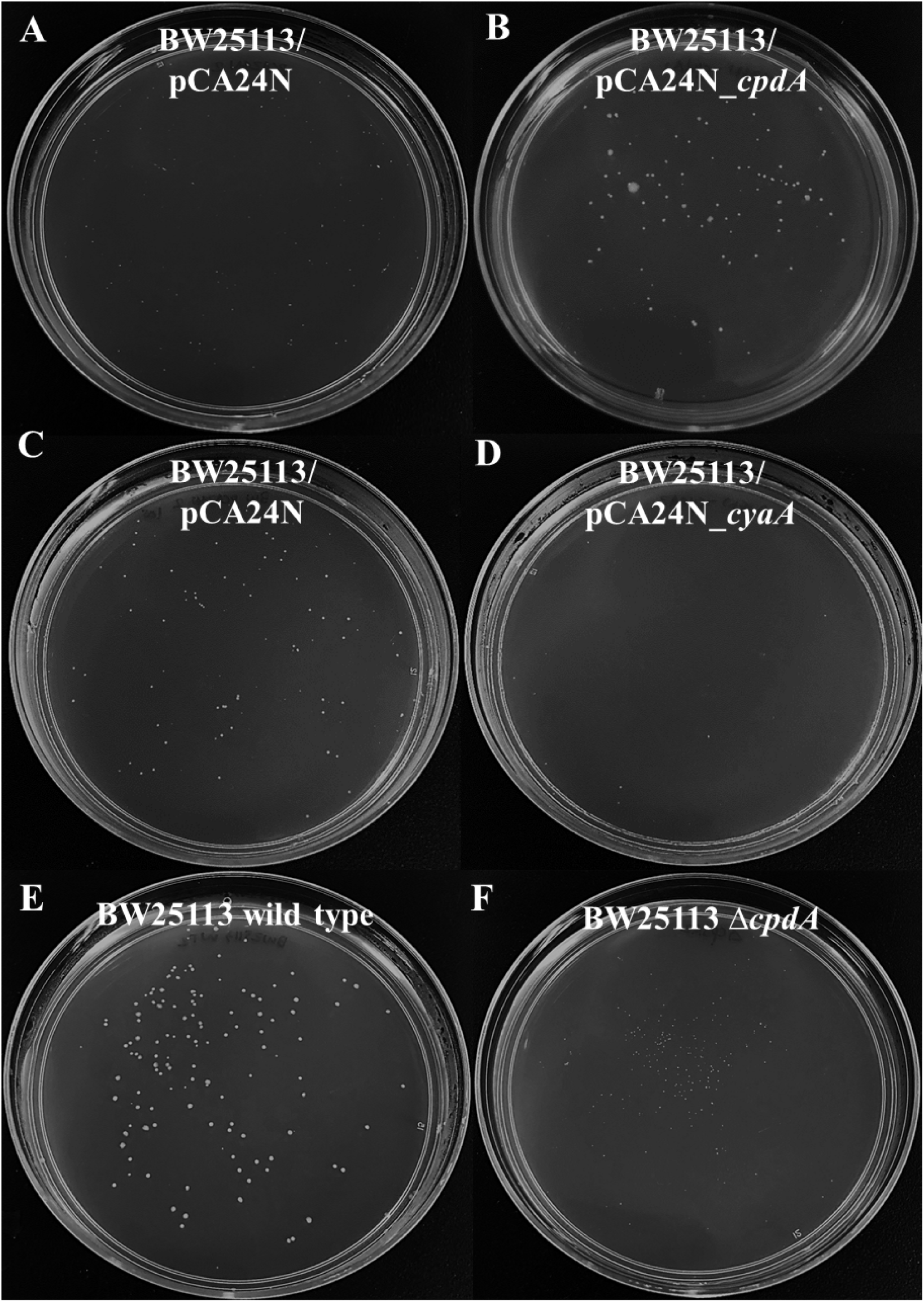
Persister cell waking on Ala agar plates after inactivating or producing proteins related cAMP. *E. coli* BW25113 persister cells with (**A**) pCA24N after 4 days, (**B**) pCA24N_*cpdA* after 4 days, **(C)** pCA24N after 5 days, **(D)** pCA24N_*cyaA* after 5 days, (**E**) wild type BW25113 after 3 days, and **(F)** BW25113 Δ*cpdA* after 3days. Plates were incubated at 37 °C and contain M9 5X Ala. One representative plate of two independent cultures is shown.

**Supplementary Figure 13.**
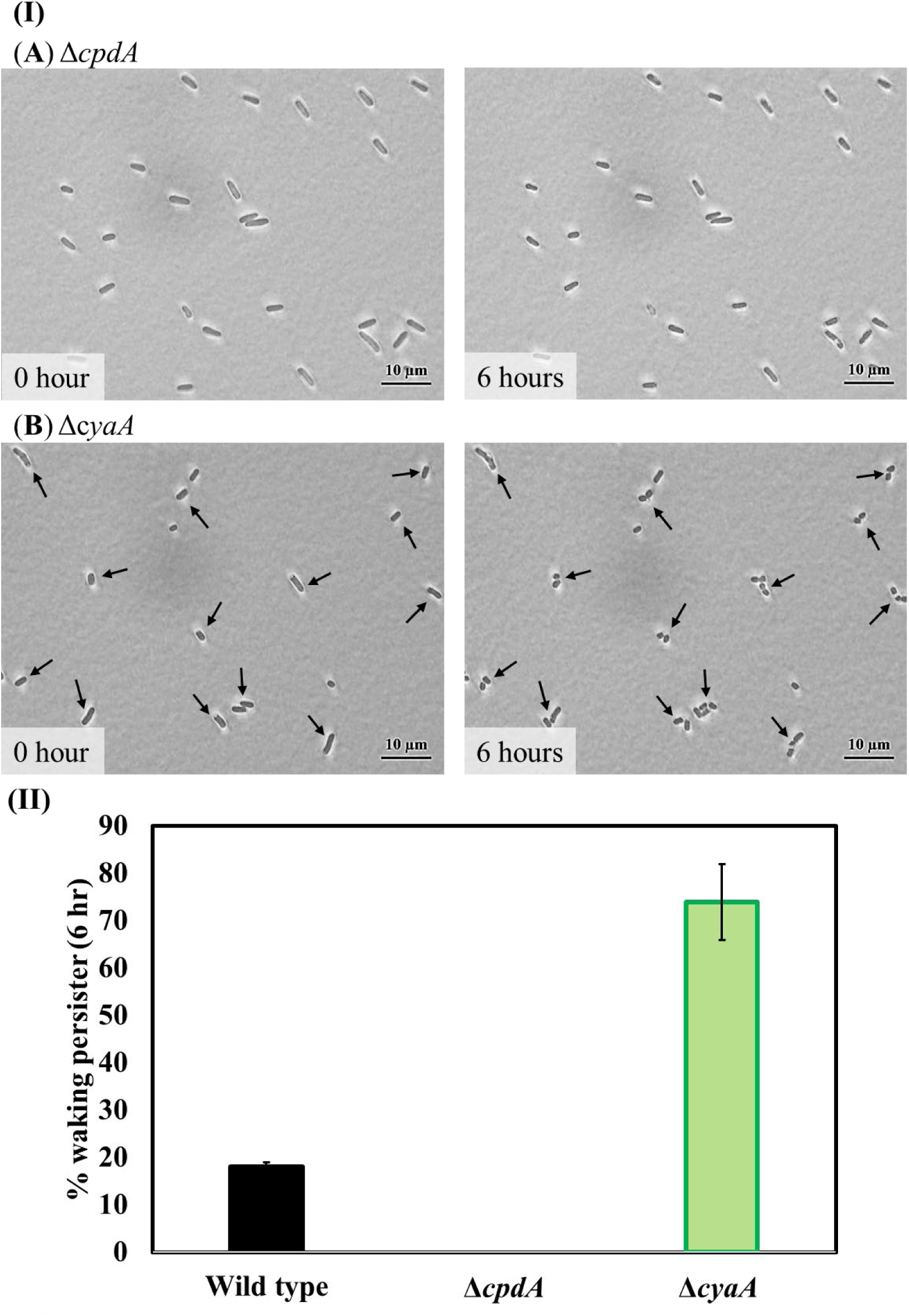
Single persister cell waking on Ala after inactivating cAMP-related proteins CpdA and CyaA. (**I**) Persister cells of (**A**) BW25113 Δ*cpdA* and (**B**) BW25113 Δ*cyaA* waking on M9 5X Ala agarose gel pads after 6 h at 37°C. Scale bar indicates 10 µm. Representative results from two independent cultures are shown. (**II**) Persister cell waking (%) after 6 hours.

**Supplementary Figure 14.**
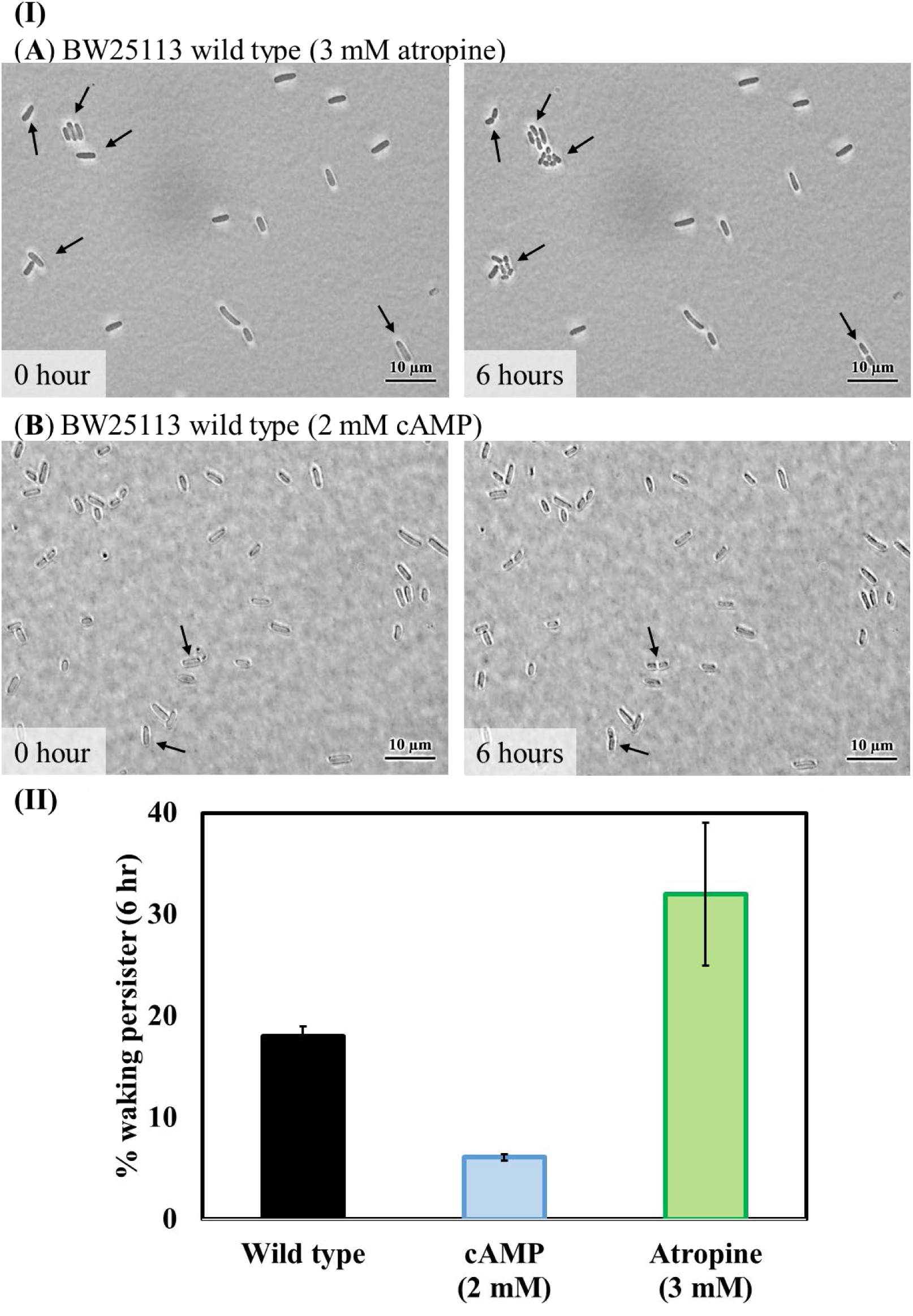
Single persister cell waking on Ala with exogenous atropine and cAMP. (**I**) Persister cells of wild-type BW25113 waking on M9 5X Ala on agarose gel pads with (**A**) 3 mM atropine or (**B**) 2 mM cAMP after 6 h at 37°C. Black arrows indicate cells that resuscitate. Scale bar indicates 10 µm. Representative results from two independent cultures are shown. (**II**) Persister cell waking (%) after 6 hours.

**Supplementary Figure 15.**
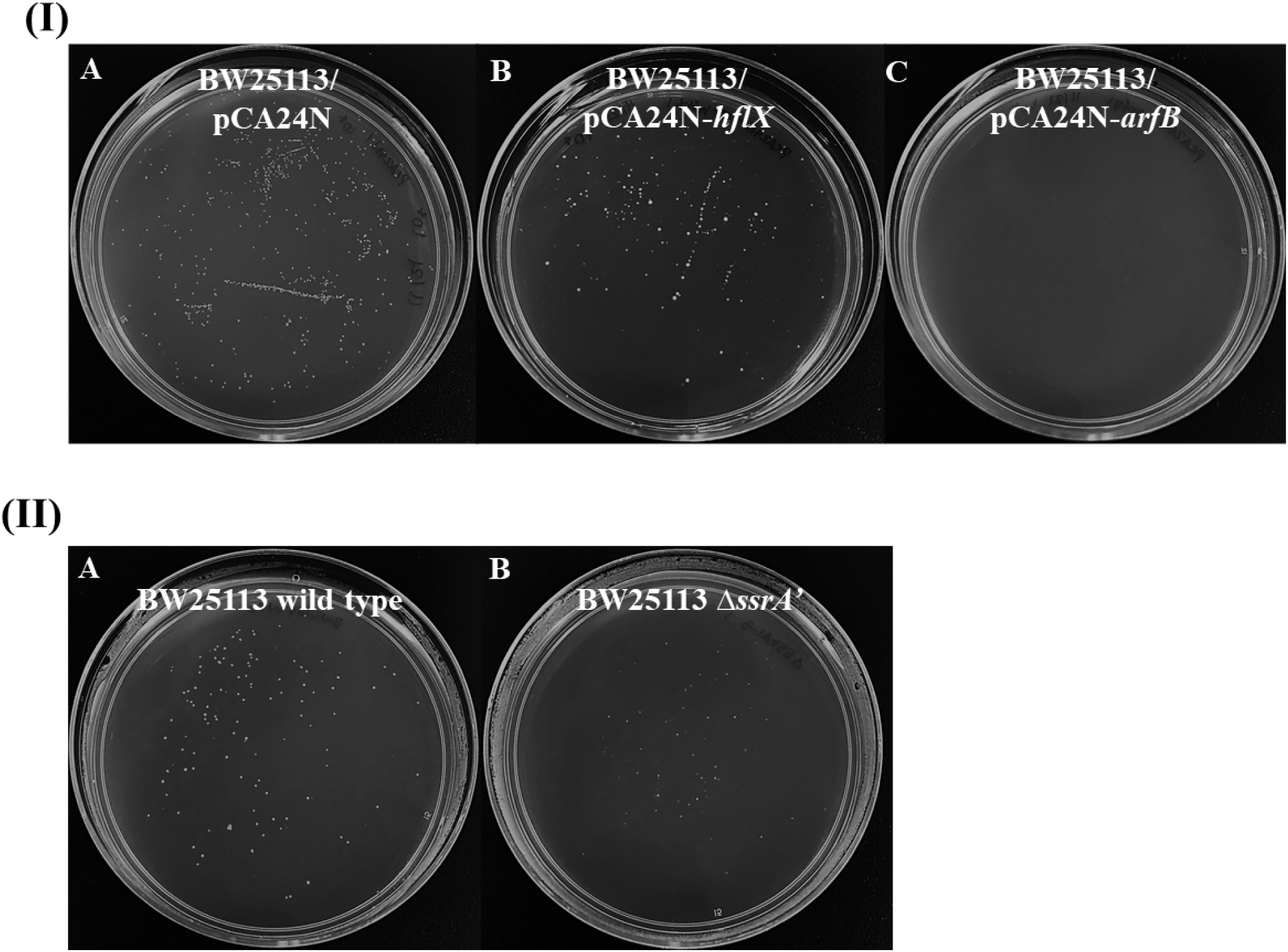
Persister cells waking on Ala agar plates after producing HflX and ArfB and inactivating SsrA. (**I**) *E. coli* BW25113 persister cells with (**A**) empty plasmid, (**B**) pCA24N_*hflX*, and (**C**) pCA24N_*arfB* were incubated at 37 °C on M9 5X Ala agar plates for 6 days. (**II**) *E. coli* BW25113 persister cells with (**A**) wild type and (**B**) BW25113 Δ*ssrA’*. One representative plate of two independent cultures is shown.

**Supplementary Figure 16.**
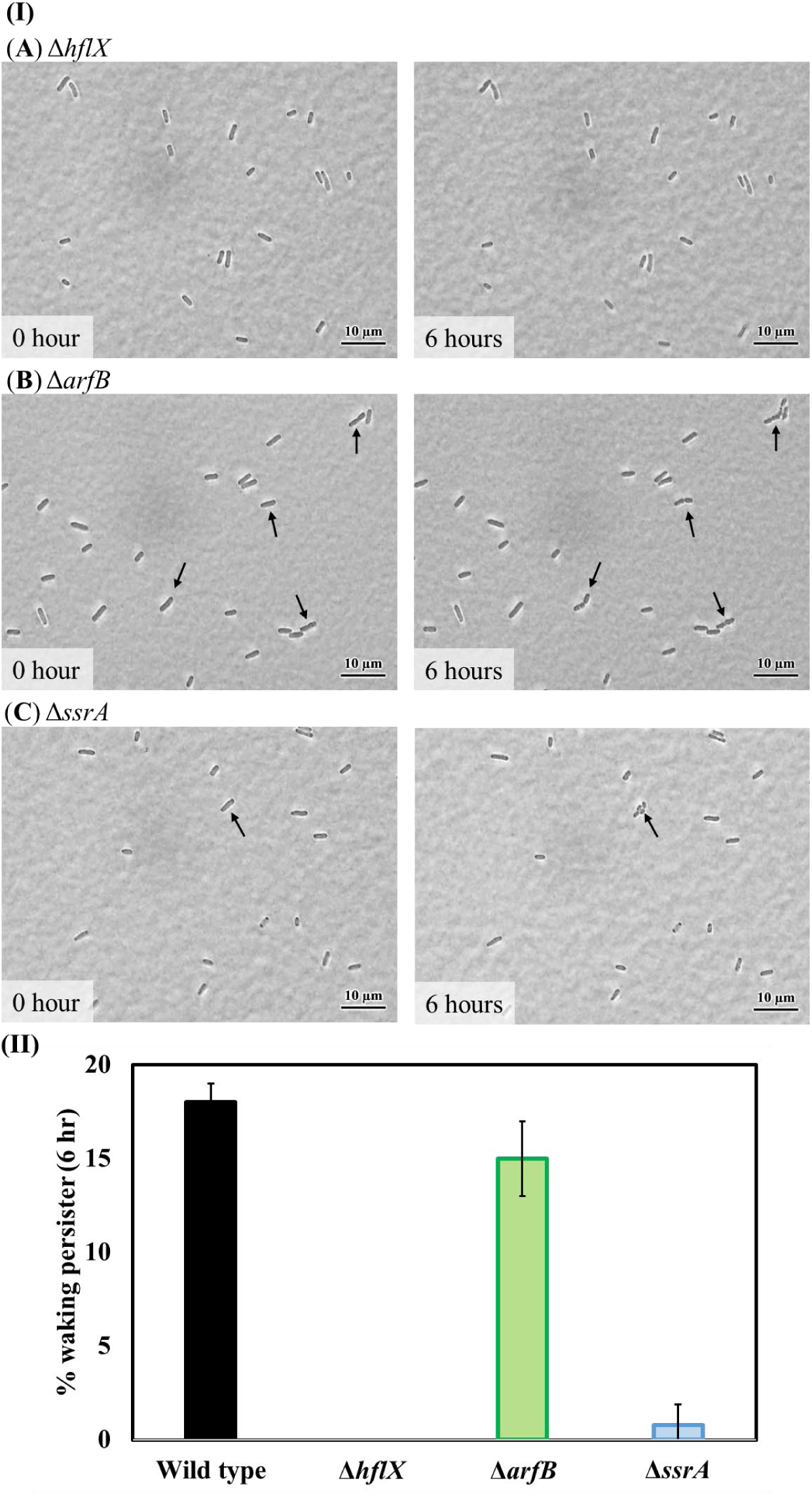
Single persister cell waking on Ala after inactivating ribosome rescue proteins. (**I**) Persister cells of (**A**) BW25113 Δ*hflX*, (**B**) BW25113 Δ*arfB*, and (**C**) BW25113 Δ*ssrA*, waking on M9 5X Ala agarose gel pads after 6 h at 37°C. Black arrows indicate cells that resuscitate. Scale bar indicates 10 µm. Representative results from two independent cultures are shown. (**II**) Persister cell waking (%) after 6 hours.

**Supplementary Figure 17.**
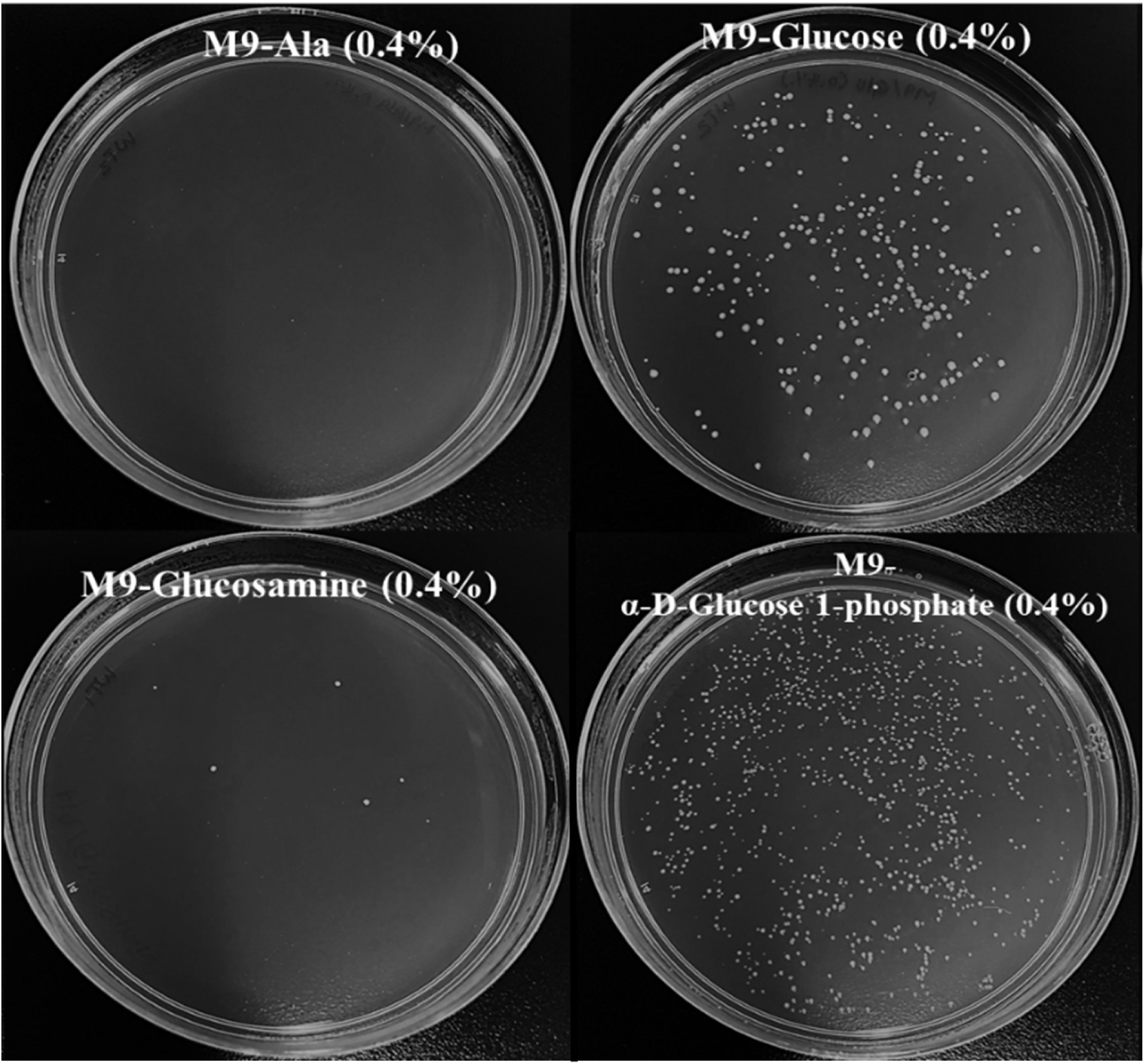
Persister cell waking on agar plates with polysaccharides as the sole carbon source. Persister cell waking of the wild type on M9 agar plates with alanine, glucose, glucosamine (D-glucosamine and N-acetyl-D-glucosamine), and α-D-glucose 1-phosphate (0.4%). Plates were incubated at 37 °C for 3 days. One representative plate of two independent cultures is shown.

**Supplementary Figure 18.**
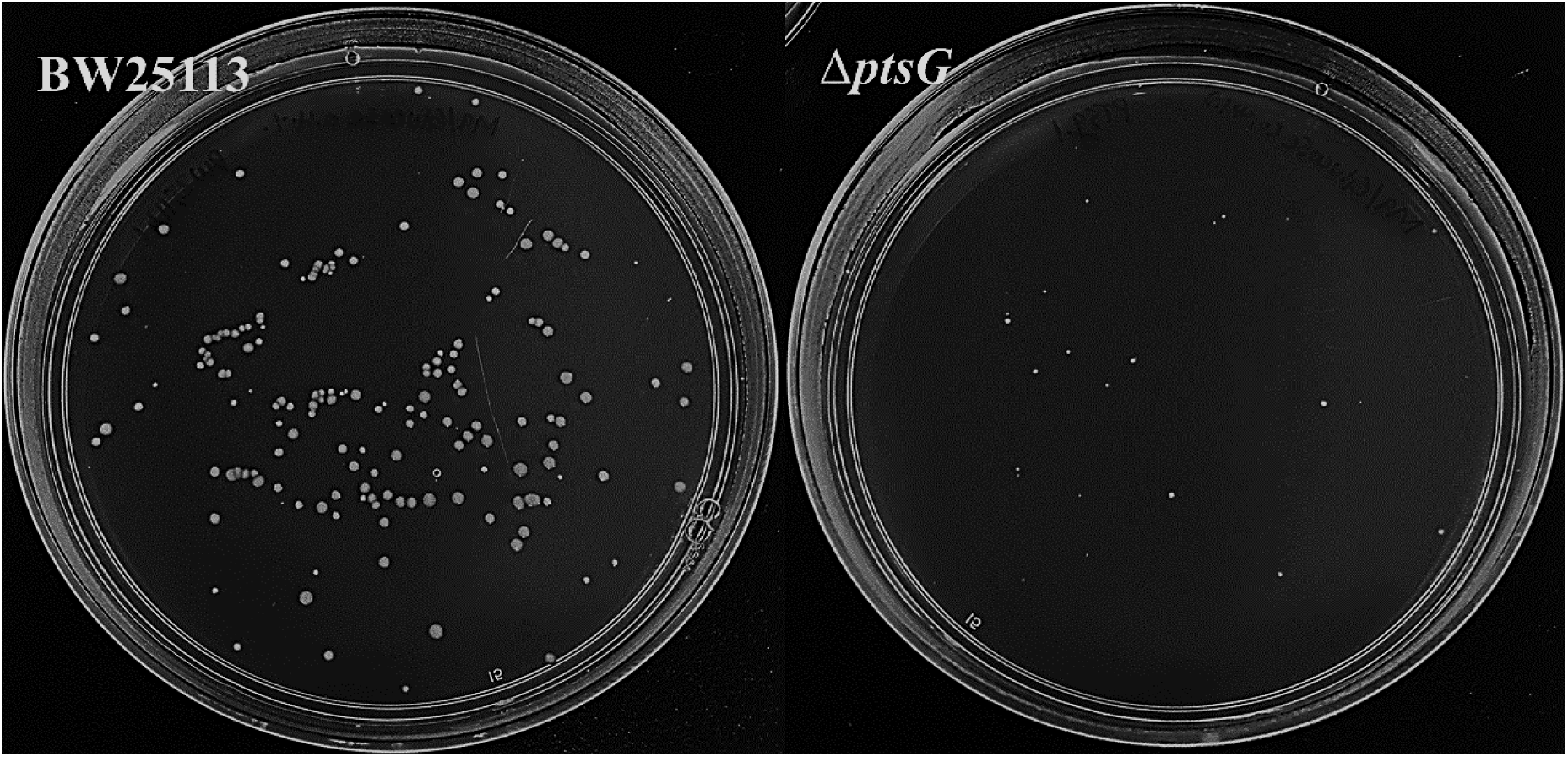
Persister cell waking on glucose agar plates after inactivating PtsG. Persister cell waking of the wild type (left panel) and the Δ*ptsG* mutant (right panel) on M9 0.4% glucose agar plates incubated at 37 °C for 2 days. One representative plate of two independent cultures is shown.

**Supplementary Figure 19.**
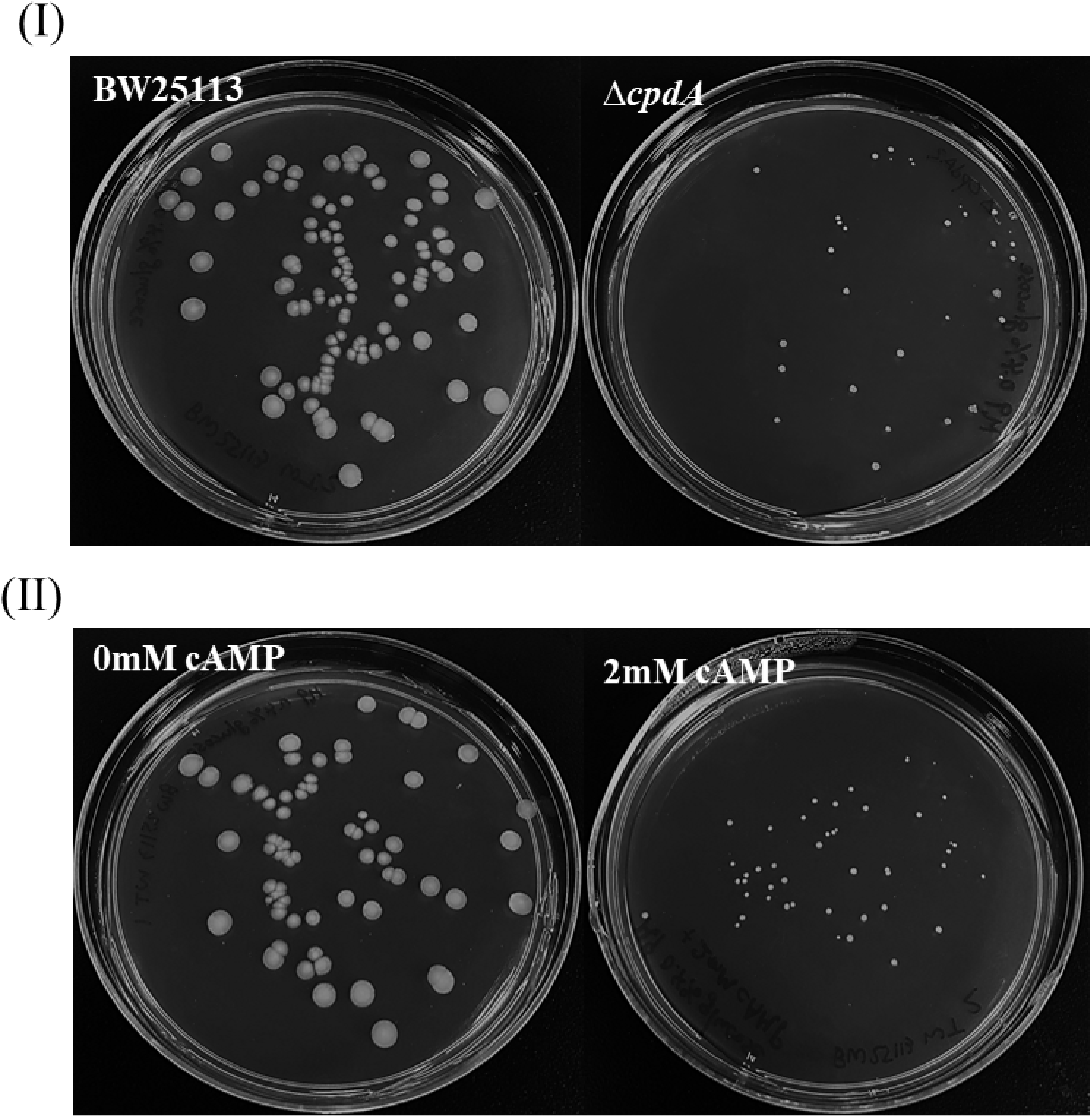
Persister cell waking on glucose agar plates after increasing cAMP. Persister cell waking of (**I**) wild type and the Δ*cpdA* mutant and (**II**) wild type with 0 mM and 2 mM cAMP on M9 0.4% glucose agar plates incubated at 37 °C on for 2 days. One representative plate of two independent cultures is shown.

**Supplementary Figure 20.**
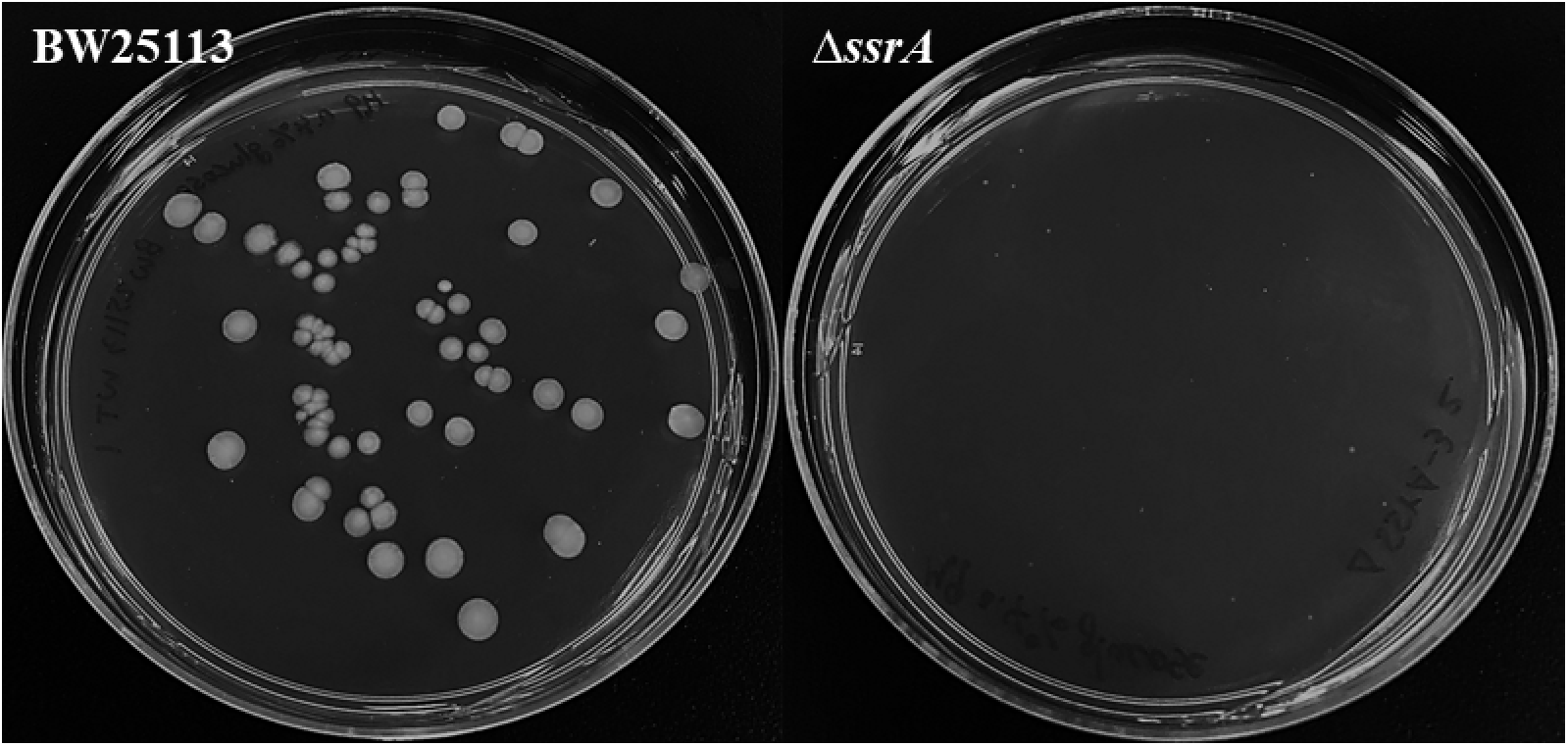
Persister cell waking on glucose agar plates for Δ*ssrA*. Persister cell waking for the wild type (left panel) and the Δ*ssrA* mutant (right panel) on M9 0.4% glucose agar plates incubated at 37 °C for 2 days. One representative plate of two independent cultures is shown.

**Supplemental video 1. Persister cell resuscitation in a glucose gradient leads to chemotaxis.** The video shows the resuscitation of wild type, Δ*cyaA*, and Δ*crr* persister cells in motility agar with a glucose gradient (0 to 5%, with the glucose at high concentrations on the left-hand side). One representative video of two independent cultures is shown.

